# Unmethylated Regions Encompass The Functional Space Within The Maize Genome

**DOI:** 10.1101/2021.04.21.425900

**Authors:** William Ricci

**Affiliations:** University of Georgia, Dept. of Plant Biology

## Abstract

Delineating the functional space within genomes has been a long-standing goal shared among geneticists, molecular biologists, and genome scientists. The genome of *Zea mays* (maize) has served as a model for locating functional elements within the gene-distal intergenic space. A recent development has been the discovery and use of accessible chromatin as a proxy for functional regulatory elements. However, the idea has recently arisen that DNA methylation data could supplement the use of accessible chromatin data for homing in on regulatory regions. Here, I test the robustness of using DNA methylation as a proxy for functional space. I find that CHG methylation can be non-arbitrarily partitioned into hypo-methylated and hyper-methylated regions. Hypo-methylated CHG regions are stable across development and contain nearly all accessible chromatin. Note: changes that will be made in version 2: expand introduction; expand discussion; add additional analyses; expand methods; link to github scripts.

## Introduction

Accessible chromatin regions (ACRs) are regions either depleted of nucleosomes or containing unstable nucleosomes, thereby allowing access to non-nucleosome DNA-binding proteins. The first survey of accessible chromatin in maize was performed with differential MNase treatment^1^ and revealed elevated accessibility at KNOTTED1 binding sites, conserved non-coding sequences, and the transcription start sites of expressed genes. Later studies performed with differential MNase^2^, DNase-seq^3, 4^, and ATAC-seq^5, 6^ collectively revealed that accessible chromatin was the probable location of cis-regulatory elements (CREs). Accumulating genetic evidence suggested that large plant genomes—most notably that of maize—contained substantial quantities of CREs within the gene-distal space, dozens of kilobases away from the nearest genes^7–11^. Researchers have attempted to home in on these remote putative CREs by narrowing the focus to ACRs. Previous genome-wide surveys of maize ACRs have yielded contrasting results. Surveys have reported 126,992^2^, 21,445^3^, 35,822^4^, and 32,111^5^ ACRs in shoot tissues and 89,455^2^ and 36,467^4^ ACRs in root. Two reports of MNase^2^ and DNase^3^ experiments were similar in both reporting ∼ 11 mb of accessible chromatin. Reports of tissue-specific ACRs included congruent estimates of 80% of ACRs shared between V2 inner stem and husk^3^ and ∼80% of leaf ACRs shared with ear primordia^5^. Slightly less congruent estimates include 23% of leaf ACR bp shared with root^2^, and ∼37% of leaf ACRs shared with root^4^. Overall, the maize ACR literature suggests that accessible chromatin is dynamic among tissues, or at a minimum, ACR estimates are sensitive to differences in experimental and computational methods. These factors pose a challenge for exhaustively surveying the regulatory space of maize. ACRs present only in rare cell types or induced only by specific environmental cues could be invisible to most experimental designs. Furthermore, weakly accessible regions or very small ACRs (for example, cryptic accessible chromatin within H3K27me3 domains^12^) are expected to fall below the detection thresholds of most methods.

In a recently published comment, Crisp et al.^13^ proposed using DNA methylation, specifically in the CHG sequence context (hereafter called mCHG), to overcome the limitations associated with accessible chromatin data. Regions with depleted mCHG levels could act as a guide to home in on the functional space within large plant genomes. The proposed benefits of using mCHG included the following:

1. Methylation cleanly partitions into hypo-methylated and hyper-methylated regions.
2. Hypo-methylated regions remain stable, despite developmental changes and environmental stressors.
3. Hypo-methylated regions encompass all ACRs, regardless of the ACRs’ developmental and stress-responsive dynamics.
4. Collectively, hypo-methylated regions provide clarity to demarcating the regulatory space, including distal CREs and the upstream edges of genes’ promoters. Such demarcations have previously remained ambiguous.
5. Furthermore, hypo-methylated regions can distinguish genuine genes from pseudogenes.

The maize methylome literature lends credence to these claims; however, as of now there has been only one study^22^ that has rigorously tested them in the context of a large plant genome. The potential utility of using DNA methylation to demarcate the functional space behooves us to test the robustness of these claims. Therefore, this manuscript aims to test the following hypotheses:

1. The maize methylome is bimodal and can be separated into clear hypo-methylated and hyper-methylated regions.
2. Hyper-methylated regions are stable across development.
3. The totality of accessible chromatin is encompassed by hypo-methylated regions.

## Results

### The Maize Methylome is Bimodal

In this section of the manuscript, I aim to achieve the following goals:

1. Test if bimodal methylation distributions persist across diverse tissue types.
2. Characterize the strength of methylation bimodality.
3. Establish a threshold for identifying discrete hypo-methylated regions in a non-arbitrary manner.

16 B73 methylomes, derived from various developmental stages, were downloaded from the NCBI Sequence Read Archive (sup. table 3.1). Some methylomes comprised multiple biological replicates, while others comprised only single samples. For consistency, replicates were merged (sup. table 3.2). The B73 AGPv4 reference genome^14^ was used for all genomic analyses in this manuscript. Chloroplast methylomes were first analyzed for quality control. Chloroplast chromatin, which is depleted of DNA methylation, was used to measure non-conversion rates—a metric that reflects the efficiency of bisulfite conversion and the severity of other artifacts that arise experimentally or *in-silico*. The weighted methylation levels (in all sequence contexts, i.e. CG, CHG, or CHH) of 100 bp non-overlapping windows on the AGPv4 chloroplast genome are shown in figure 3.1A. For most of the methylomes, the non-conversion rates approached zero, with some skew up to 1 or 2 percent (fig. 3.1A). Coleoptile and endosperm (9 DAP) skewed higher than did other samples but rarely exceeded 5% non-conversion (fig. 3.1F). The major outlier was ear (silking), in which a proportion of 0.19 windows exceeded 5% non-conversion (fig. 3.1F). In summary, all methylomes—with the exception of ear (silking)— displayed acceptable non-conversion rates, unlikely to interfere with subsequent analyses.

**Figure 3.1:**
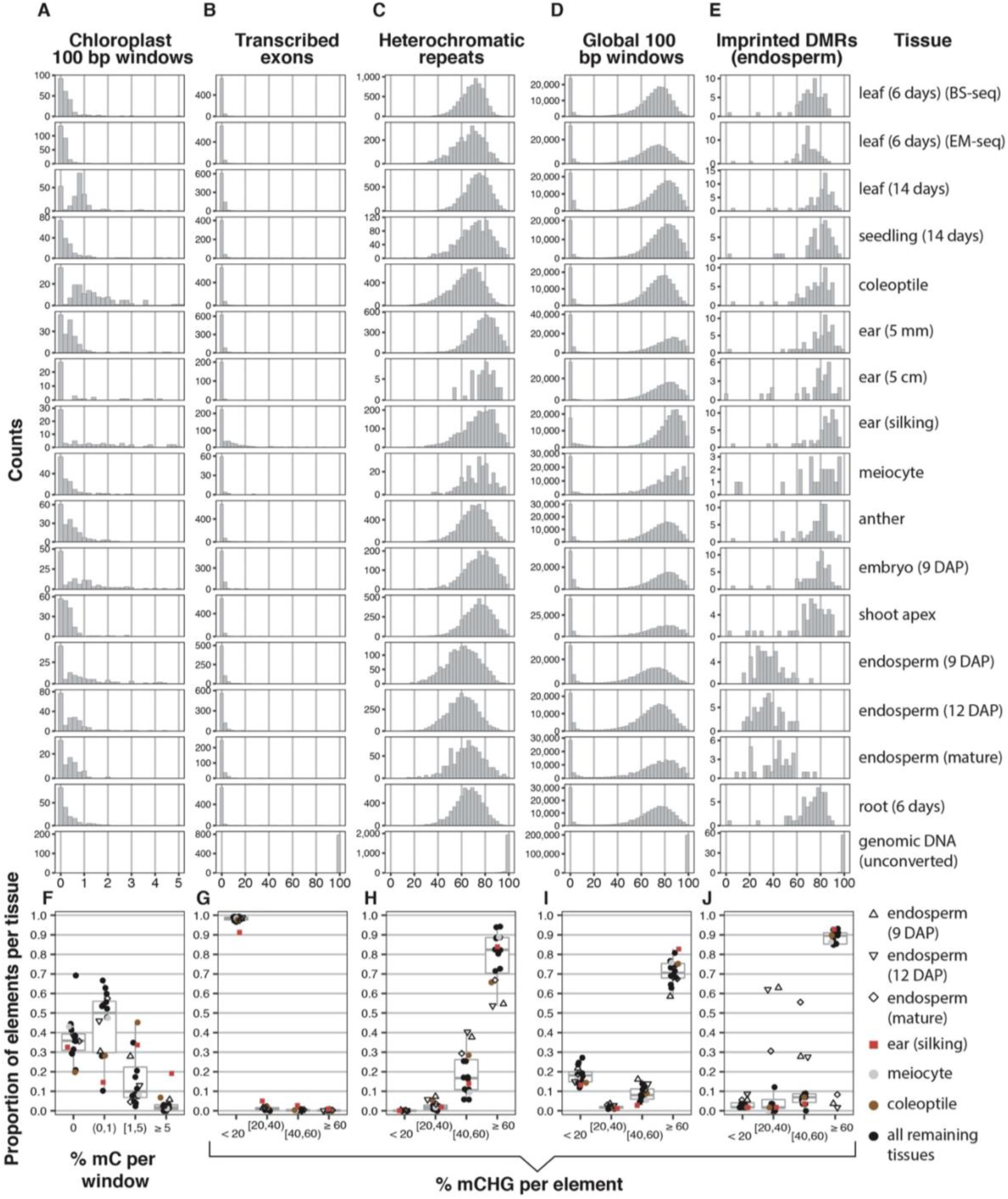
The Global Distribution of CHG Methylation. Only elements meeting minimum mapping threshold were analyzed. Element numbers vary per methylome, depending on how many meet mapping threshold. **A-E**: Frequency histograms of % methylation for different types of elements. Each row corresponds to a different methylome (denoted in rightmost column). **F-J**: The proportion of elements falling into discrete % methylation bins. Each point corresponds to a single methylome (denoted on right). Boxplots indicate quartiles. The genomic DNA (unconverted) sample was omitted. **A & F**: % mC (i.e. all sequence contexts) for 100 bp windows in the reference chloroplast sequence. This column serves to illustrate non-conversion rates. **B & G:** % mCHG for constitutively transcribed exons. This column serves as a positive control for developmentally stable, hypo-methylated regions. **C & H:** % mCHG for knob180 and TR-1 heterochomatic repeats. This column serves as a positive control for developmentally stable, hyper-methylated regions. **D & I:** % mCHG for 100 bp non-overlapping windows. 200,000 windows were randomly selected from each methylome. **E & J:** % mCHG for endosperm-imprinted DMRs. CHG-specific DMRs selected from Table S5 of Zhang et al. (2014). This column serves as a positive control for regions that vary in methylation status among homologous chromosomes.

I next identified positive control regions that consistently held hypo- or hyper-methylated (in the CHG context) across all methylomes. Exons within constitutively highly expressed genes (sup. data file 1) were used for the hypo-methylated control. Such exons were expected to constitutively exclude mCHG. figure 3.1B shows the % mCHG at exons in all 16 methylomes. Nearly all exons were at or approaching 0% mCHG. Among all methylomes, a median proportion of 0.99 exons fell below 20% mCHG (fig. 1G). Ear (silking) was the only outlier, with a proportion of 0.91 exons below 20% mCHG (fig. 3.1G). A parallel analysis of mCG (sup. fig. 3.1B and G) revealed a large proportion of hyper-methylated exons (median proportion of 0.61 exons < 20% mCG), which was expected because of the known prevalence of gene body methylation^15^. For the constitutively hyper-methylated control, I used knob180 and TR-1 repeats (sup. data file 2), the main repetitive elements comprising the heterochromatic knobs. Despite variation in mapping sparseness, the heterochromatic repeats were CHG- and CG-hyper-methylated in all methylomes (fig. 3.1C and sup. fig. 3.1C). Mean % mCHG ranged from approximately 60% to 80% across methylomes and mean % mCG was > 80% in all methylomes. Although the mean % mCHG exceeded 60% in all methylomes, the tails of many distributions dipped below 40% mCHG (fig. 3.1C and H). Importantly, virtually none of the heterochromatic repeats dipped below 20 % mCHG or mCG (median proportion 0.0004 in mCHG and 0 in mCG) (fig. 3.1H and sup. fig. 3.1H). In summary, the controls satisfied the expectations of constitutive hypo- or hyper-methylation, indicating that the methylomes were technically comparable.

I next calculated the methylation of 100 bp non-overlapping windows, genome-wide. Both mCHG and mCG displayed clear bimodalities in all methylomes, wherein 100 bp windows partitioned into discrete hypo-methylated (< 20%) and hyper-methylated (> 40%) modes (fig. 3.1D and sup. fig. 3.1D). Mid-methylated windows (20-40%) were strongly depleted, with median proportions of 0.017 for mCHG (fig. 3.1I) and 0.0056 for mCG (sup. fig. 3.1I). One could imagine several biological scenarios that weaken or interfere with methylome bimodality:

1. windows containing mixtures of methylated and unmethylated cytosines in similar number
2. small hypo-methylated regions within windows, causing resolution artifacts
3. differential methylation at the same window on homologous chromosomes
4. differential methylation among cells within a sample
5. differential methylation among pooled plants

The strong depletion of mid-methylated windows suggested that all of these scenarios were rare or non-existent. For certainty, I included a positive control of regions that were differentially methylated (DMRs) between maternal and paternal chromosomes in endosperm (fig. 3.1E and 3.1J, sup. fig. 3.1E and 3.1J) (DMRs from Zhang et al., 2014^16^; sup. data files 3 and 4). In the three endosperm methylomes (9 DAP, 12 DAP, and mature), the DMRs ranged from approximately 20-60% mCHG and mCG, with modes centered around 40%. If chromosome-specific DMRs were prevalent, modes between 20-50% should have been visible in global methylation distributions, which they were not, even in endosperm (although endosperm methylomes did show slight increases in mid-methylation bins (fig. 3.1I)). This indicated that among all methylomes surveyed, methylation status was largely consistent among loci of homologous chromosomes.

Although the mid-methylated windows are rare, it is necessary to explain the small proportion that do exist. These windows could be transitions from hypo-to hyper-methylated regions, and thus should reside within close proximity to hypo-methylated windows. Alternatively, the windows could be standalone and independent of hypo-methylated regions or allele-specific hypo-methylated regions. To address this question, I plotted the mid-methylated windows’ distances to the nearest hypo-methylated windows. Figures 3.2A and 3.2B show these plots from a representative methylome, leaf (6 days old). It was evident that most of the mid-methylated windows resided in close proximity to hypo-methylated windows, effectively acting as transitions between hypo- and hyper-methylated regions. The proximities from all methylomes are quantified in figures 3.2C and D, which display the proportions of mid-methylated windows directly adjacent to or within 1 kb of hypo-methylated windows. As methylation approached 20%, mid-methylated windows were increasingly in close proximity to hypo-methylated windows. Meiocyte displayed anomalous low proportions that could be attributed to the high sparseness of its methylome. Approximately 77% of mCHG and 93% of mCG mid-methylated windows could be explained by proximity to hypo-methylated windows. The unexplained mid-methylated windows may be standalone; however, it was unknown how many of these windows could be attributed to stochastic methylation variation. Collectively, the evidence suggested that mid-methylated regions did not constitute independent elements.

**Figure 3.2:**
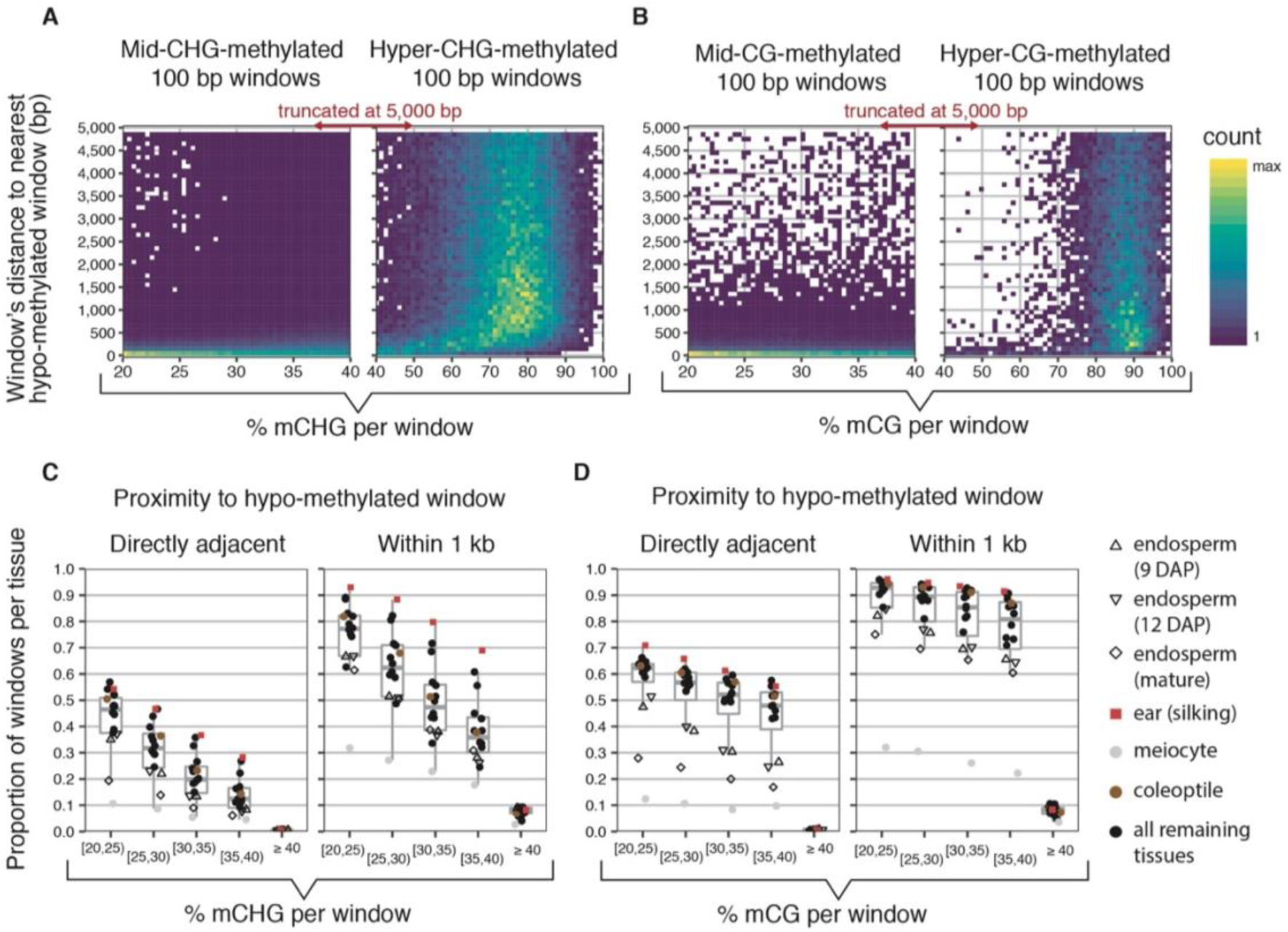
The distance of mid-methylated windows to the nearest hypo-methylated windows. The % methylation of mid-methylated (20-40%) and hyper-methylated (> 40%) 100 bp windows—in both the CHG and CG sequence contexts—are plotted against their distances to the nearest hypo-methylated (< 20 %) window. For each sequence context, random hyper-methylated windows were selected to match the number of hypo-methylated windows. For mCHG there are 164,337 windows and for mCG there are 35,457 windows. **A & B:** The leaf (6 days) (BS-seq) methylome is shown as a representative sample. Plotted are 2D histograms of % methylation versus distance to nearest hypo-methylated window in the CHG context (**A**) and CG context (**B**). **C & D:** The proportion of 100 bp windows falling into discrete % methylation bins that satisfy the criteria of being either adjacent to or within one kb of the nearest hypo-methylated window. All methylomes shown. Each point corresponds to a single methylome (denoted on right). Boxplots indicate quartiles.

In summary, there is little compelling evidence to refute the bimodal methylome. 20% mCHG and mCG acts as a non-arbitrary threshold for identifying hypo-methylated regions. Supplementary figure 3.2 shows parallel analyses performed in the CHH sequence context. mCHH remained consistently low across all elements and was unsuitable for identifying hypo-methylated regions. Therefore, mCG and mCHG were used to identify regions that were either lowly methylated (< 20%) or lacking methylation (hereafter known as unmethylated regions—UMRs).

### Unmethylated Regions (UMRs) are Stable Across Development

This section assesses the stability of UMRs across diverse developmental stages. UMRs were identified with a simple algorithm, starting with 100 bp windows sliding in 20 bp increments. Windows were required to overlap at least 20 reads and at least four informative cytosines. Windows with weighted methylations below 20% mCHG or mCHG/mCG were retained, merged, and edge-refined. UMRs shorter than 50 bp were discarded. UMR sets were created for all methylomes. Additionally, a Mo17 methylome was mapped to B73 to serve as a positive control with the expectation of moderate UMR instability—prior literature has identified many differentially methylated regions among the B73 and Mo17 inbred lines^17^.

Descriptive overviews of the UMR sets can be found in supplementary table 3.2 and supplementary figures 3.3-3.8. The raw UMR sets are available in the supplementary data files. UMRs shorter than 150 bp displayed elevated methylation levels and variation, probably attributable to the lower coverage, cytosine counts, and increased inherent stochasticity of small UMRs (sup. fig. 3.3). Consequently, in later analyses, UMRs were separated by length. Generally, genic UMRs (i.e. UMRs overlapping genes) were four-fold larger than distal UMRs (> 5 kb from genes) (sup. fig. 3.6). The lengths of UMRs and total UMR counts varied substantially among methylomes (sup. fig. 3.4 and sup. fig. 3.6). This variation could be attributed to UMR fragmentation caused by methylome sparseness: UMR counts anticorrelated with coverage and UMR length, while coverage positively correlated with UMR length (sup. fig. 3.7). As a quality control, UMRs were compared to distal enhancers defined by Oka et al. (2017) (sup. fig. 3.8). Virtually all enhancers were recapitulated by the UMRs sets, with the exception of Mo17 and meiocyte.

The varying UMR counts, UMR lengths, and library coverages posed a challenge for UMR comparison among methylomes. Therefore, a combinatorial approach was used in which the UMR set from each methylome was superimposed onto the other methylomes. Figures 3.3A and B show the results of UMRs 150 bp and larger. Each row corresponds to a methylome and its primary UMR set. Points within plots represent the proportion of primary UMRs, when superimposed onto a secondary methylome, that fall within a given % mCHG bin. Values in the < 20% mCHG bin represent the proportions of primary UMRs that were conserved as UMRs in other methylomes. The ‘transcribed exons’ column displays only UMRs overlapping constitutively expressed exons. These UMRs are expected to be conserved among all B73 methylomes. For example, the leaf (6 days) (BS-seq) exon UMRs (fig. 3.3A, top left panel), when superimposed onto the Mo17 methylome (denoted by red x), showed a UMR conservation of 0.94. In other words, 6% of these UMRs did not remain hypo-methylated in the Mo17 methylome. In contrast, the B73 methylomes, with the exception of ear (silking), showed nearly complete UMR conservation. With the exception of ear (silking), the exon UMR control held true virtually 100% of the time for all methylomes.

**Figure 3.3:**
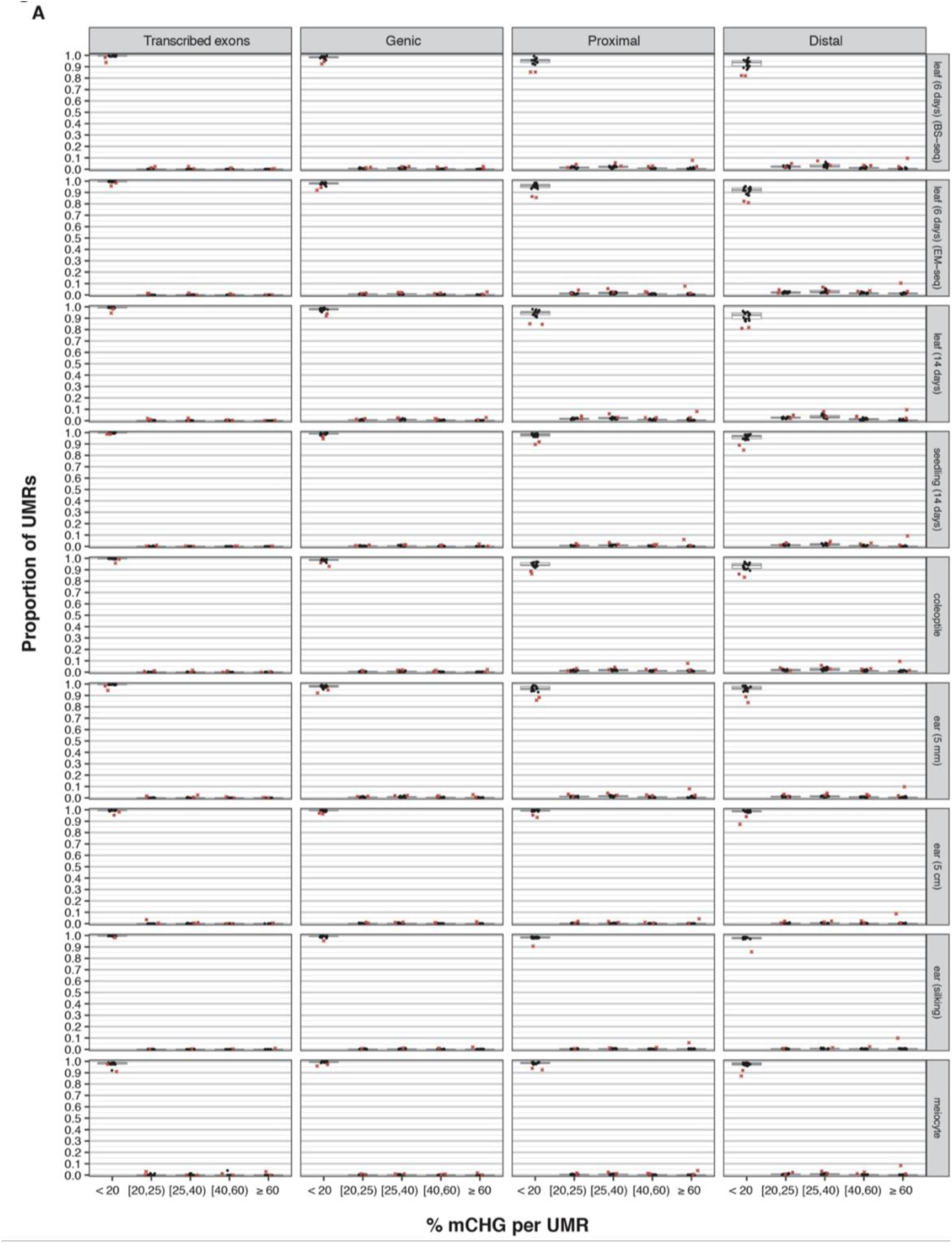
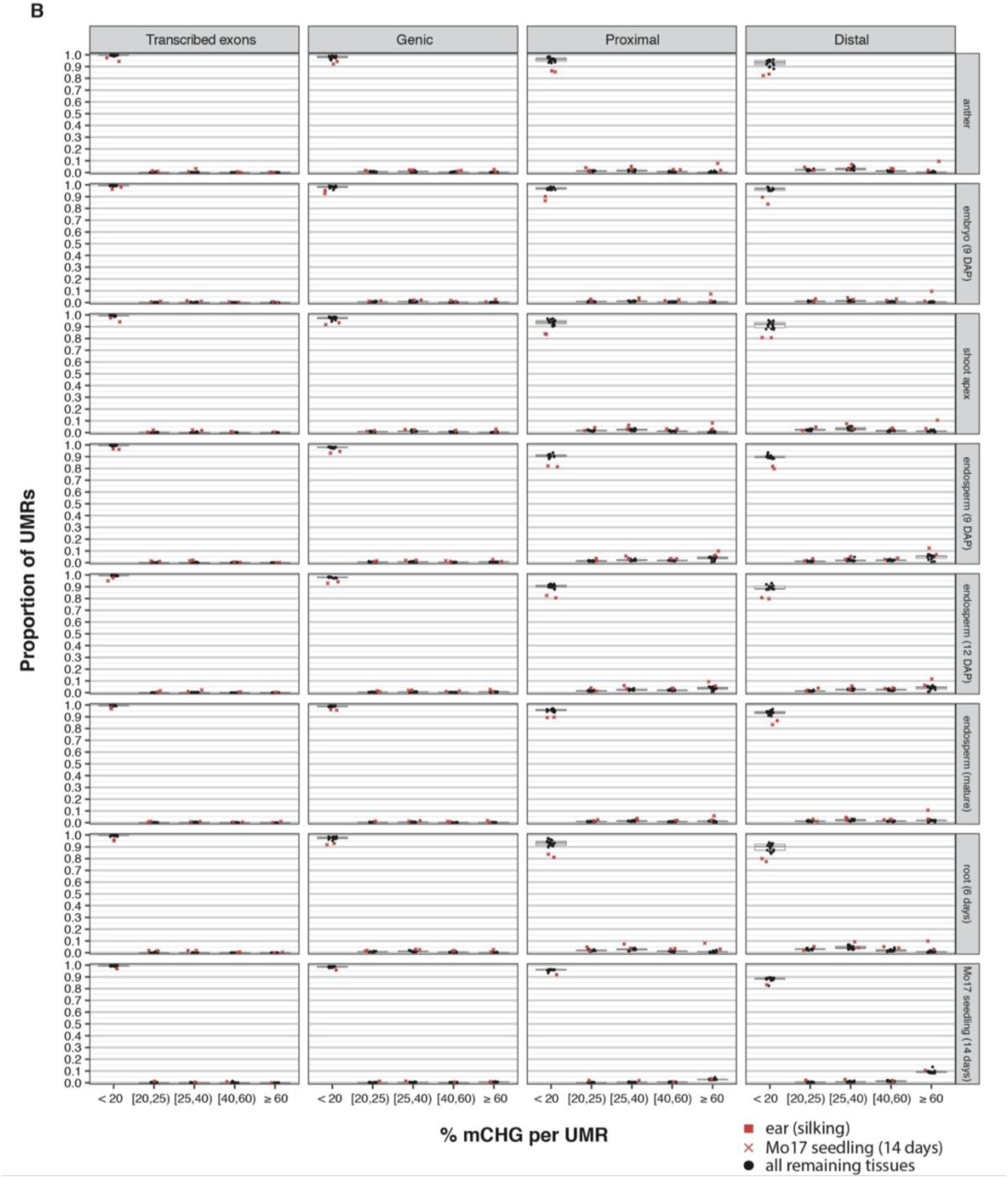
UMR (> 150 bp) stabilities across methylomes. Shown are the proportions of a tissue’s UMRs falling into discrete % mCHG bins in other tissues. Each row corresponds to a set of UMRs (> 150 bp in length) identified in the primary methylome indicated on the right. Points within plots indicate the proportions of methylation levels in secondary methylomes. Boxplots indicate median and quartiles. Primary tissue UMRs are split by their proximities to AGPv4 annotated genes. Genic: overlaps genes by at least 1 bp. Proximal: within 5,000 bp of genes. Distal: > 5,000 bp from genes. Also shown are primary tissue UMRs that overlap constitutively transcribed exons. These serve as positive controls for developmentally stable UMRs. Ear (silking) and Mo17 seedling (14 days) are differentiated by shape and color (indicated in legend). **A & B:** Identical formatting but different primary methylomes.

Genic UMRs were the most conserved, while distal UMRs were the least conserved. For example, in leaf (6 days) (BS-seq), median UMR conservation was 0.98 (genic), 0.95 (proximal), and 0.94 (distal). Median proportions of distal UMR conservation ranged from 0.99 (ear (5 cm)) to 0.88 (endosperm (12 DAP)). UMRs shorter than 150 bp showed considerably less conservation across methylomes (sup. fig. 3.9) than did UMRs longer than 150 bp. Furthermore, breaking down UMRs into 100 bp intervals revealed a relationship between UMR length and stability that could explain the elevated conservation of genic UMRs. Supplementary figure 3.10A shows the median proportions of conserved UMRs from all methylomes, broken down by UMR length. Supplementary figure 3.10B shows pairwise comparisons among methylomes of biologically similar tissues or biological replicates; these comparisons provide values for baseline reproducibility. The baseline-corrected stabilities (sup. fig. 3.10C) of small UMRs (e.g. 50-250 bp) were highly variable. However, the stabilities of UMRs larger than 350 bp exceeded 0.90, and with increasing length, approached complete conservation. The length-stability curves were similar for genic, proximal, and distal UMRs, which suggested that the greater lengths of genic UMRs could explain their greater conservation. UMRs from methylomes comprising merged biological replicates were moderately more stable than UMRs from single biological samples (sup. fig. 3.11). This was expected, considering that merged replicates should conceal nonreproducible UMRs. However, the length-stability curves persisted, suggesting that experimental variation alone could not fully account for the reduced stability of small UMRs. B73 methylomes were ranked by the baseline-corrected stabilities of their distal UMRs (sup. fig. 3.12). Root (6 days) ranked as the least stable; although, with increasing UMR length, it converged with the other methylomes (sup. fig. 3.10C). It is noteworthy that the methylomes ranked as most stable (e.g. ear (5 cm)) also had relatively low library coverages, suggesting that low coverages could artificially inflate UMR stabilities.

An analysis of non-conserved UMR methylation revealed that non-conserved UMRs generally did not become hyper-methylated (sup. fig. 3.13). With the exception of endosperm and Mo17 UMRs, non-conserved UMRs remained near 20% mCHG and were depleted in the hyper-methylated mode, a trend that strengthened with increasing UMR length (compare sup. fig. 3.13A, B, and C). In contrast, non-conserved endosperm UMRs frequently became hyper-methylated (> 60% mCHG). The 9 DAP, 12 DAP, and mature endosperm non-conserved UMRs (less than 500 bp) were hyper-methylated in 46.43%, 39.78%, and 29.94% of cases, respectively. UMRs larger than 500 bp were not enriched for hyper-methylation. Overall, as much as ∼5% of total endosperm UMRs may switch to a hyper-methylated state in a different tissue.

In summary, the vast majority of UMRs are stable across development; however, UMRs smaller than ∼300 bp should be handled with caution, especially among endosperm tissues. UMRs larger than 500 bp can be considered developmentally static. For practical purposes, a consensus UMR set was created by merging the reciprocally overlapping UMRs from three high-coverage methylomes (available in sup. data files). This consensus UMR set was used in most analyses hereafter.

### UMRs Encompass Accessible Chromatin Regions

I next sought to test if accessible chromatin regions (ACRs) were fully contained within ACRs, regardless of the developmental dynamics of the accessibility. I used publicly available ATAC-seq, DNase, and differential MNase datasets, derived from diverse tissue types (sup. table 3.3). Genrich (https://github.com/jsh58/Genrich) was used to identify ACRs, and based on the peak-calling efficacy, 8 tissues were kept for downstream analysis: earshoot (5 mm), earshoot (1 cm), leaf (6 days old), coleoptilar node, 1mm root tip, mesophyll, inner husk, and inner stem. Quantifying the degree of ACR overlap with transcription start site positive controls revealed high false-negative rates in all tissues (sup. table 3.4). This suggested that analyses based on ACR peak calls would not be exhaustive.

In order to identify developmentally dynamic ACRs, all tissues were compared to each other in pairwise to identify overlapping ACRs. ACRs were grouped into categories from 0 to 7, based on the number of tissues containing overlapping ACRs, with 0 denoting a tissue-specific ACR and 7 denoting an ACR that overlaps other ACRs in every tissue. These ACR categories were tested for overlap with UMRs (fig. 3.4). ACRs were compared against the consensus UMR set, as well as leaf (6 days) and root (6 days) UMR sets filtered by lengths of 150 bp and larger. With an overlap threshold of 1 bp, the vast majority of ACRs overlapped UMRs (fig 3.4A). Category 2-7 ACRs exceeded a 99% UMR overlap rate for genic, proximal, and distal ACRs. However, the overlap of tissue-specific ACRs (category 0) dropped substantially, down to approximately 97% (genic), 93% (proximal), and 92% (distal). This suggested that as much as 8% of distal ACRs resided outside of UMRs. Alternatively, an 8% false-positive distal peak-calling rate or false negative UMR-calling rate could yield a similar effect. The steep shift between category 0 and 1 ACRs suggested that the necessity of replication for category 1 could reduce false-positive peak calls.

**Figure 3.4:**
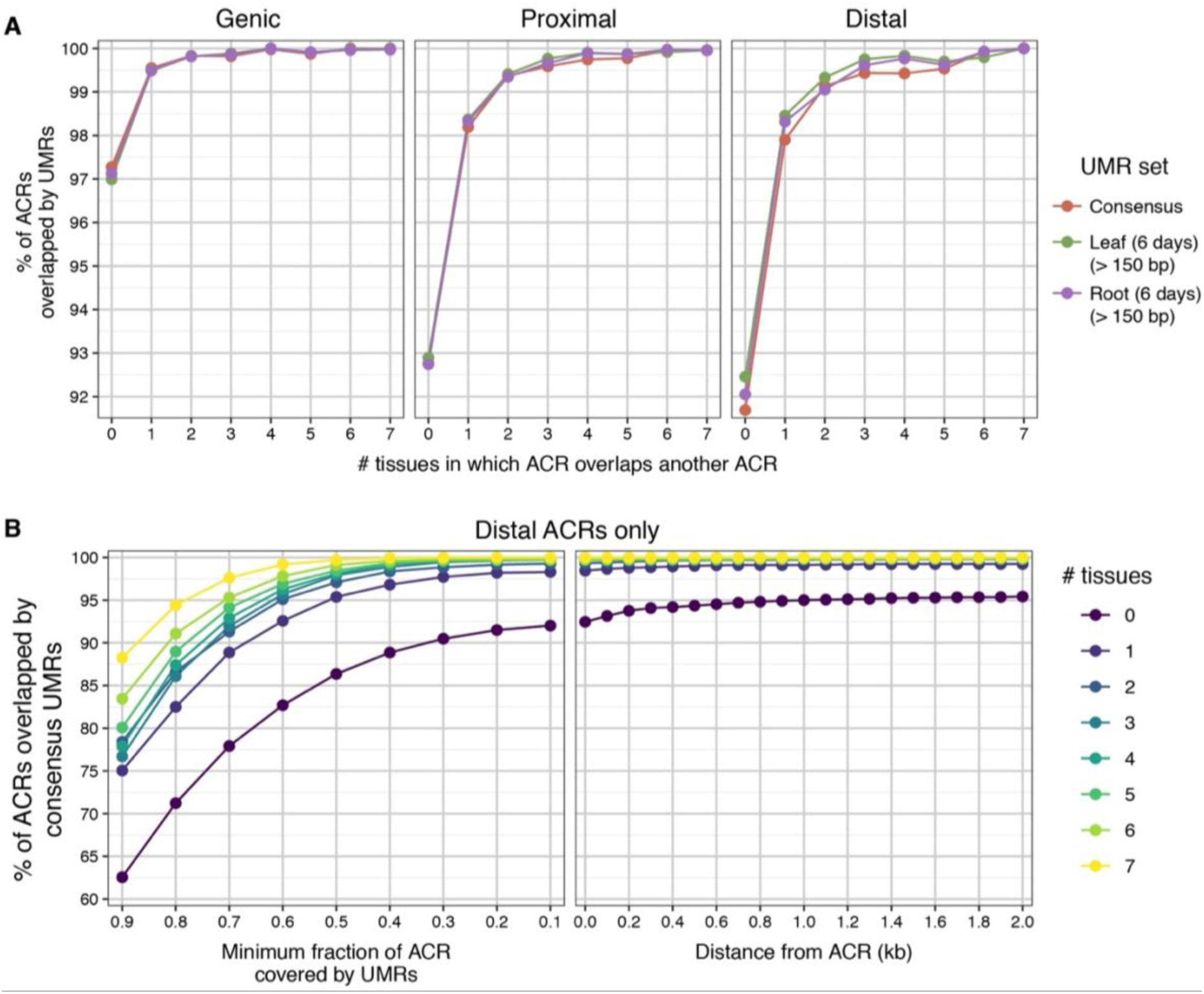
ACR and UMR overlap frequencies. Shown here are overlaps of UMRs with accessible chromatin regions (ACRs). ACR peaks were called in 8 different tissues. ACRs were categorized based on how many ACRs from other tissues they overlapped. Category 0 ACRs do not overlap any other ACRs. Category 7 ACRs overlap ACRs from all 7 other tissues. **A:** The % of ACRs from each category that overlap UMRs by ≥ 1 bp. ACRs are separated by gene proximity. **B:** Gene-distal ACRs are shown only. Shown in left panel are the % of ACRs with different amounts of overlap with UMRs. For example, ∼86% of category 0 ACRs are covered by at least 50% of their lengths with UMRs. Shown in the right panel are the % of ACRs that have a UMR within the specified distance.

The nature of the ACR-UMR overlaps were analyzed in greater depth in fig. 3.4B, which breaks down the fractions of ACRs overlapping UMRs (for distal ACRs only). More than 95% of category 1-7 ACRs were contained within UMRs by at least half of their ACR lengths. However, the percent overlap decreased as the fraction approached complete overlap, suggesting that as much as 1/4 of distal category 1-7 ACRs overlapped UMRs but simultaneously overlapped space outside of UMRs. The remainder of ACRs were fully contained within UMRs. To account for artifacts associated with peak-calling edges, ACRs were expanded outward by 2 kb in order to detect UMRs within close proximity. This expansion increased the category 0 ACR overlap rate from 92% to 95%, leaving a remaining 5% of ACRs that were isolated within the intergenic space. Analogous analyses for genic and proximal ACRs are shown in sup. fig. 3.14. Proximal ACRs displayed trends similar to distal ACRs, while genic ACRs showed even greater UMR overlap.

While nearly all ACRs were contained within UMRs, a large number of UMRs did not overlap ACRs in any of the 8 tissues (sup. fig. 3.15): 15,739 (distal), 9,260 (proximal), and 7,149 (genic) UMRs were ACR-free. The high false-negative rate of ACR calls had the potential to inflate these counts. To deal with the artifacts associated with a qualitative peak-calling approach, I quantified the ATAC-seq and DNase-seq read coverage over UMRs (sup. fig. 3.16). ACR-free UMRs still showed coverage elevated above that of the distal UMR negative control regions. This suggested that weakly accessible chromatin, undetectable by the peak caller, may exist within a subset of the ACR-free UMRs. Therefore, the ACR-free UMRs were split into quartiles of coverage and these quartiles were analyzed as separate UMR categories hereafter.

I next sought to test whether the dynamic ACRs and ACR-free UMRs displayed genetic evidence of functionality. The high-density of genotyped SNPs among maize lines, coupled with the rapid decay of linkage disequilibrium (∼2 kb), allows for kilobase-resolution mapping of quantitative trait loci^18^. Therefore, I tested UMRs for genetic associations with gene expression and agronomic traits. Strong-effect QTLs linked to 41 agronomic traits^7^ and eQTL corresponding to the expression variation of ∼28,000 genes^19^ were used for the analysis. UMRs falling within 2 kb of QTL polymorphisms (to account for linkage disequilibrium) were counted as QTL hits. Overall, all but the bottom quartile of ACR-free UMRs (Q1) showed significantly elevated frequencies of QTL hits, compared to the background rate of the negative control regions (fig. 3.5). The large sample size of the Q1 UMR set (n = 3,855), along the high p-value of 0.45 (hypergeometric test), indicated a distinct absence of QTL enrichment within this UMR subset. In contrast, the three other quartiles of the ACR-free UMRs were all significantly enriched for QTL overlaps, suggesting functional enrichment in ACR-free UMRs. QTL enrichment was further increased for UMRs containing ACRs. Furthermore, enrichment increased progressively as UMRs became more constitutively accessible, but decreased at the most constitutively accessible UMRs. The input SNPs showed no overlap bias among UMR groups, indicating that the differences in QTL enrichment among groups could not be attributed to mapping artifacts.

**Figure 3.5:**
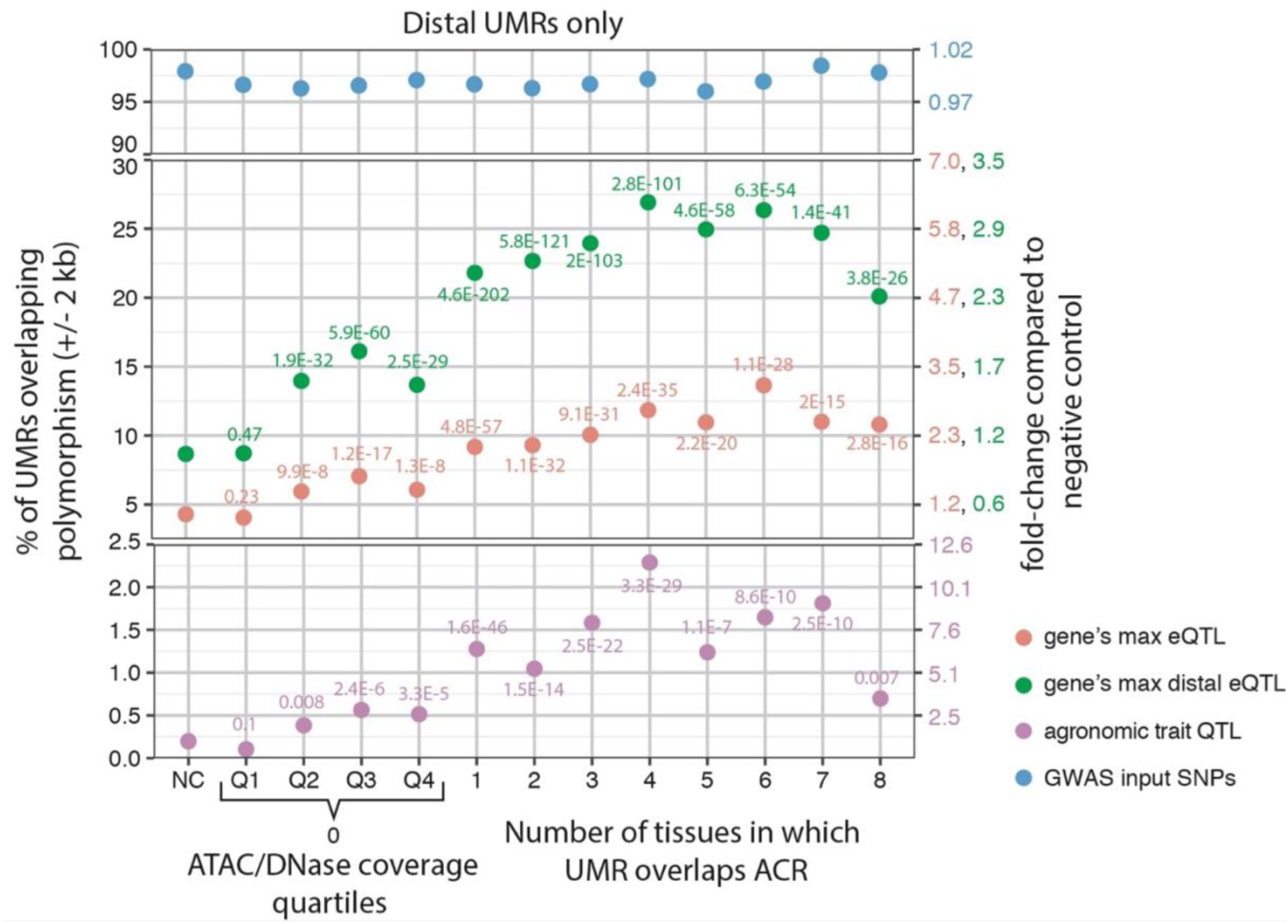
Proportions of distal UMRs overlapping QTL. Shown here are distal UMRs only. UMRs are broken down into various subcategories on the x-axis. Categories 1-8 are UMRs that overlap ACRs in 1-8 tissues, respectively. Category 0 means that UMRs overlap ACRs in no tissues. Category 0 is subdivided into quartiles based on the ATAC/DNase coverages shown in sup. fig. 3.16B. The purpose of the coverage quartiles is to account for potential false negative peak calls. Category “NC” denotes the negative control regions, which are distal regions that match the distal UMRs for count, length, and coverage. The y-axis shows the % of UMRs from each category that is within 2 kb of polymorphisms denoted in legend. Gene’s max eQTL: the global top eQTL SNP hit for each gene (measured by SNP’s r^2^ value). Gene’s max distal eQTL: the global top eQTL hit for each gene that falls > 5 kb from any annotated gene. Ag trait QTL: polymorphisms to 41 different agronomic traits, filtered for RMIP values > 10. GWAS input SNPs: the total SNPs that were used for eQTL mapping, derived from the maize HapMap 3. eQTL data were from Kremling et al. (2018) and agronomic QTL data were from Wallace et al. (2014).

It is possible that the top three quartiles of the non-ACR UMRs actually did contain ACRs, despite being missed by the peak caller. This could explain the elevated QTL hits for these quartiles compared to Q1. To address this, I relaxed the Genrich peak-calling stringency to a p-value cutoff of 0.1, which should reduce the false negative rate, although with the tradeoff of producing many false-positives. Upon relaxing the stringency, 25% of negative control regions overlapped ACRs, 30% of Q1 UMRs overlapped ACRs, and 66-76% of Q2-Q4 UMRs overlapped UMRs (sup. fig. 3.17). The disproportionate increase in the Q2-Q4 UMRs suggested the presence of cryptic accessible chromatin within these UMR classes. This left only the 3,922 UMRs within Q1 as probable ACR-free UMRs, the only group lacking genetic evidence of functional enrichment. These UMRs belonged to the consensus UMR set, which comprised reciprocally overlapping UMRs from both leaf and root, and therefore were reproducibly robust. Furthermore, the Q1 UMR set had a median length of 384 bp, with the top quarter exceeding 742 bp. As shown in the previous section, UMRs of this size are robust. Therefore, these data collectively suggest the existence of a population of UMRs lacking ACRs and lacking genetic evidence of functionality.

## Discussion

The demarcation of the genome’s functional space is a long-standing goal among plant biologists and plant breeders. ACRs have served as a useful proxy for homing in on the regulatory space in large plant genomes^3, 5^. However, the dynamic nature of ACRs and the virtually unstudied effect of stressors on ACR induction in maize means that the regulatory space may extend beyond what has been described in ACR studies. Indeed, the ∼11 megabases of ACRs previously reported^2, 3^ is far exceeded by the ∼40 megabases of UMRs in the intergenic space alone and the ∼170 megabases of UMRs in total. Furthermore, UMRs can be identified with greater confidence than can ACRs. Because UMRs fall into two virtually qualitative categories—hypo-methylated or hyper-methylated—they can be identified without subjective biases and without the need for an input mapping control. The rise of accurate long-read sequencing technology^20^, coupled with enzymatic methyl-seq^21^, will allow UMRs within the highly repetitive portion of the genome to be identified. Currently, this portion of the genome is off-limits to assays that rely on small read technology, including bisulfite-seq, ATAC-seq, DNase-seq, and MNase-seq. Therefore, the potential of UMRs may be even greater with the adoption of these new technologies.

The counts and total megabases of called UMRs varied substantially among tissues (sup. fig. 3.7). These differences could largely be attributed to variation in library sparseness. It is notable that the variation in UMR counts and median lengths decreased as methylomes increased in coverage, indicating that the higher coverage libraries stabilized and converged on similar values. The consensus UMR library was constructed out of three high-coverage libraries (leaf (Bisulfite 6 days), leaf (EM-seq 6 days), and root (6 days)) for this reason. The number of consensus UMRs identified here (95,367) was similar to the 99,422 UMRs called on the coleoptile methylome by Oka et al^3^.

The practical utility of DNA methylation behooves us to rigorously compare it to accessible chromatin data. Here, I report that nearly all ACRs are contained by UMRs. However, up to 1/4 of ACRs may spill over the edge of the UMR boundary, akin to the asymmetric placement of ACRs at the edge of transcription units. It still must be determined if the cis-regulatory elements within the ACRs spill over the UMR boundaries as well. Parsing the ACRs into the 0-7 tissue specificity metric revealed that constitutive and developmentally dynamic ACRs are similarly contained within UMRs. This indicates that the UMRs from a single tissue are sufficient to encompass the ACRs specific to a different tissue. This is further supported by the nearly identical overlap rates of the leaf, root, and consensus UMR sets. However, the exception to this is the most extreme tissue-specific ACRs, present in only 1 of the 8 tissues. As much as 8% of these ACRs may not be contained within UMRs, although that number may decrease with increasing peak-calling stringency.

Prior literature has reported enrichment of expression and agronomic QTL at intergenic ACRs^2, 5^. Rodgers-Melnick et al.^2^ reported approximately 2x enrichment of agronomic QTL hits at intergenic ACRs, compared to a negative control. Ricci, Lu, and Ji et al. ^5^ reported approximately 1.5x enrichment of eQTL at intergenic ACRs. The overlap frequencies reported here do not differ substantially; however, overlap rates are greater or weaker, depending on the UMR type. Generally, UMRs that are accessible in more tissues are more enriched for QTL, with UMRs containing ACRs from 4 or 6 tissues being the most enriched. UMRs overlapping 0 ACRs are still enriched for QTL, suggesting that there are intergenic functional elements present in the genome that would be missed by solely relying on accessible chromatin data. However, it is likely that these UMRs actually do contain ACRs that were below detection thresholds. This suggests that the vast majority of UMRs may actually contain ACRs and that ACR-free UMRs may be rare genomic elements. Collectively, these data demonstrate the utility of using UMRs to demarcate the functional space in the large maize genome.

---

**Supplementary Figure 3.1:**
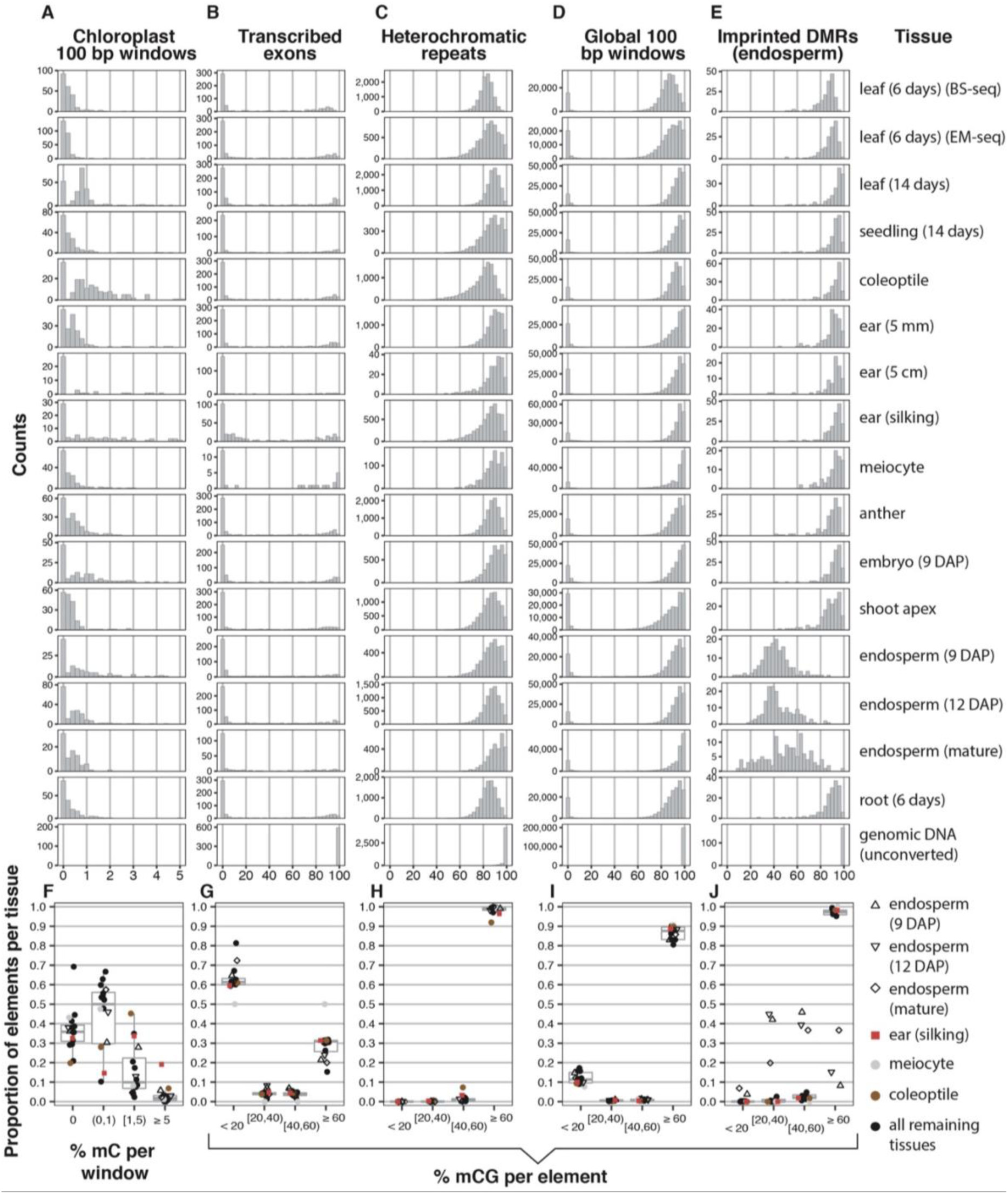
The Global Distribution of CG Methylation. Only elements meeting minimum mapping threshold were analyzed. Element numbers vary per methylome, depending on how many meet mapping threshold. **A-E**: Frequency histograms of % methylation for different types of elements. Each row corresponds to a different methylome (denoted in rightmost column). **F-J**: The proportion of elements falling into discrete % methylation bins. Each point corresponds to a single methylome (denoted on right). Boxplots indicate quartiles. The genomic DNA (unconverted) sample was omitted. **A & F**: % mC (i.e. all sequence contexts) for 100 bp windows in the reference chloroplast sequence. This column serves to illustrate non-conversion rates. **B & G:** % mCG for constitutively transcribed exons. This column serves as a positive control for developmentally stable, hypo-methylated regions. **C & H:** % mCG for knob180 and TR-1 heterochomatic repeats. This column serves as a positive control for developmentally stable, hyper-methylated regions. **D & I:** % mCG for 100 bp non-overlapping windows. 200,000 windows were randomly selected from each methylome. **E & J:** % mCG for endosperm-imprinted DMRs. CG-specific DMRs selected from Table S5 of Zhang et al. (2014). This column serves as a positive control for regions that vary in methylation status among homologous chromosomes.

**Supplementary Figure 3.2:**
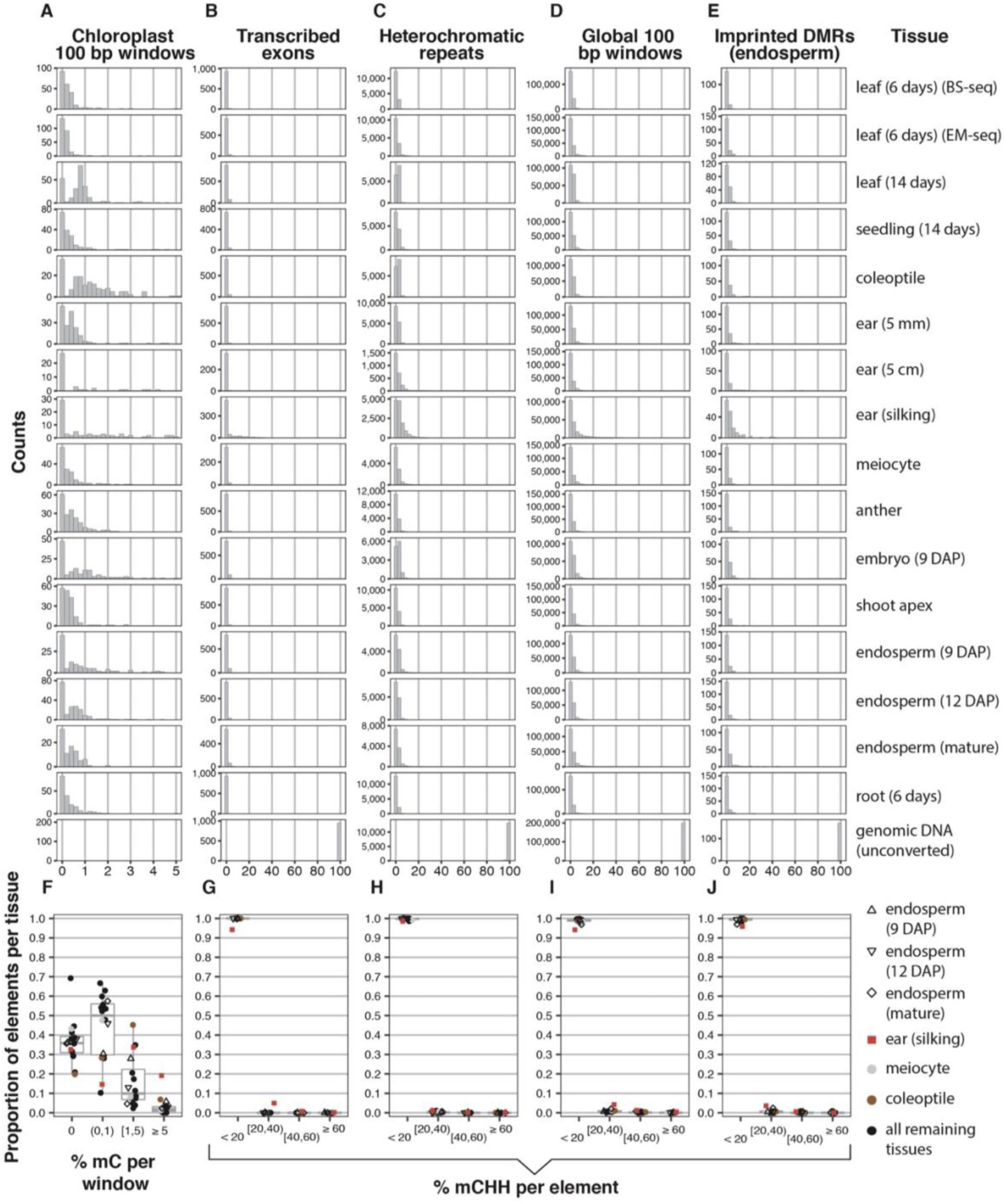
The Global Distribution of CHH Methylation. Only elements meeting minimum mapping threshold were analyzed. Element numbers vary per methylome, depending on how many meet mapping threshold. **A-E**: Frequency histograms of % methylation for different types of elements. Each row corresponds to a different methylome (denoted in rightmost column). **F-J**: The proportion of elements falling into discrete % methylation bins. Each point corresponds to a single methylome (denoted on right). Boxplots indicate quartiles. The genomic DNA (unconverted) sample was omitted. **A & F**: % mC (i.e. all sequence contexts) for 100 bp windows in the reference chloroplast sequence. This column serves to illustrate non-conversion rates. **B & G:** % mCHH for constitutively transcribed exons. This column serves as a positive control for developmentally stable, hypo-methylated regions. **C & H:** % mCHH for knob180 and TR-1 heterochomatic repeats. This column serves as a positive control for developmentally stable, hyper-methylated regions. **D & I:** % mCHH for 100 bp non-overlapping windows. 200,000 windows were randomly selected from each methylome. **E & J:** % mCHH for endosperm-imprinted DMRs. CG-specific DMRs selected from Table S5 of Zhang et al. (2014). This column serves as a positive control for regions that vary in methylation status among homologous chromosomes.

**Supplementary Figure 3.3:**
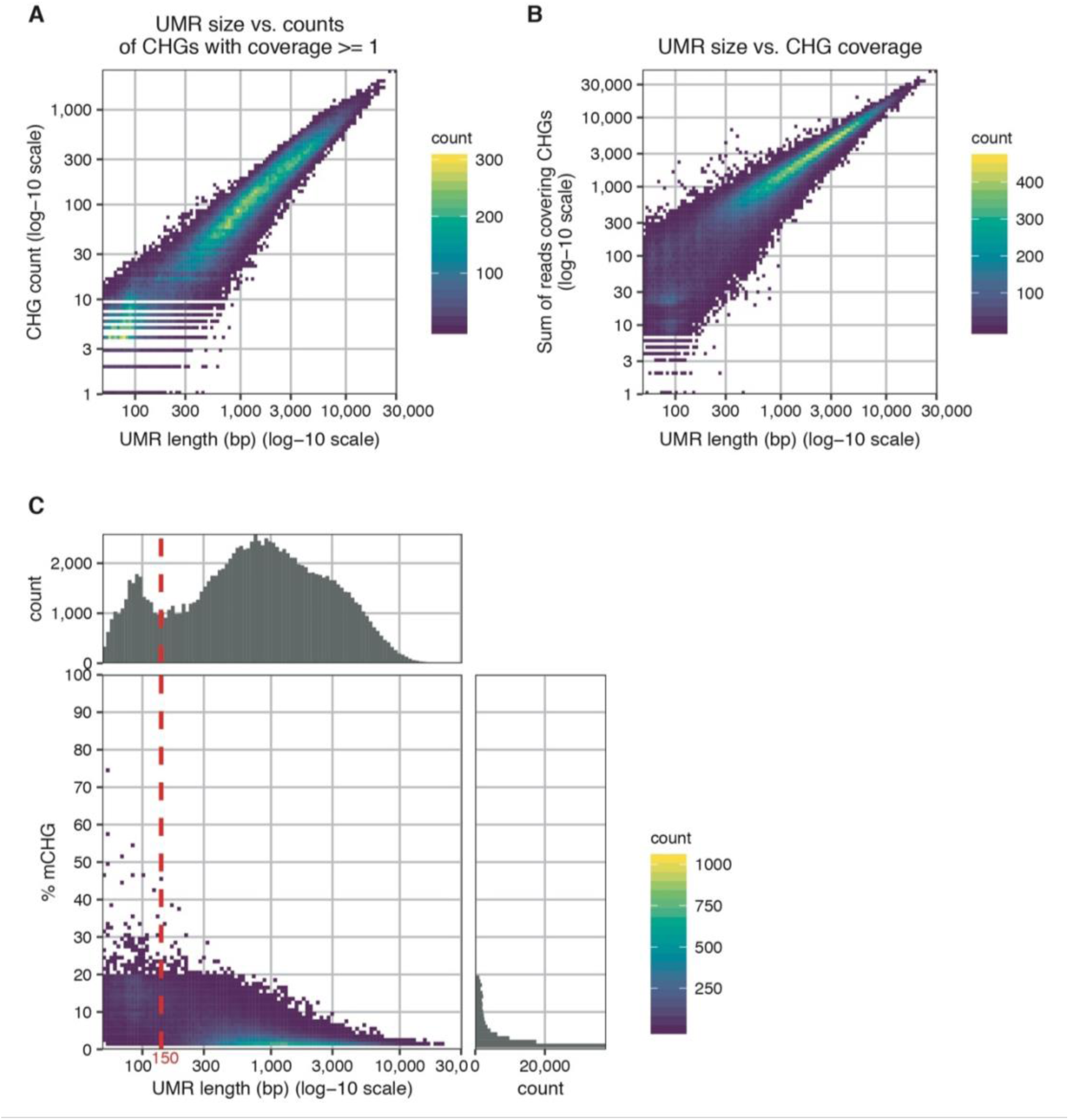
Relationships among various UMR attributes. The UMR set from leaf (6 days) (BS-seq) is shown as a representative methylome. **A:** UMR length vs. counts of covered CHGs per UMR. **B:** UMR length vs. sum of reads per UMR that cover CHGs. **C:** UMR length vs. % mCHG per UMR. Red dashed line indicates 150 bp threshold.

**Supplementary Figure 3.4:**
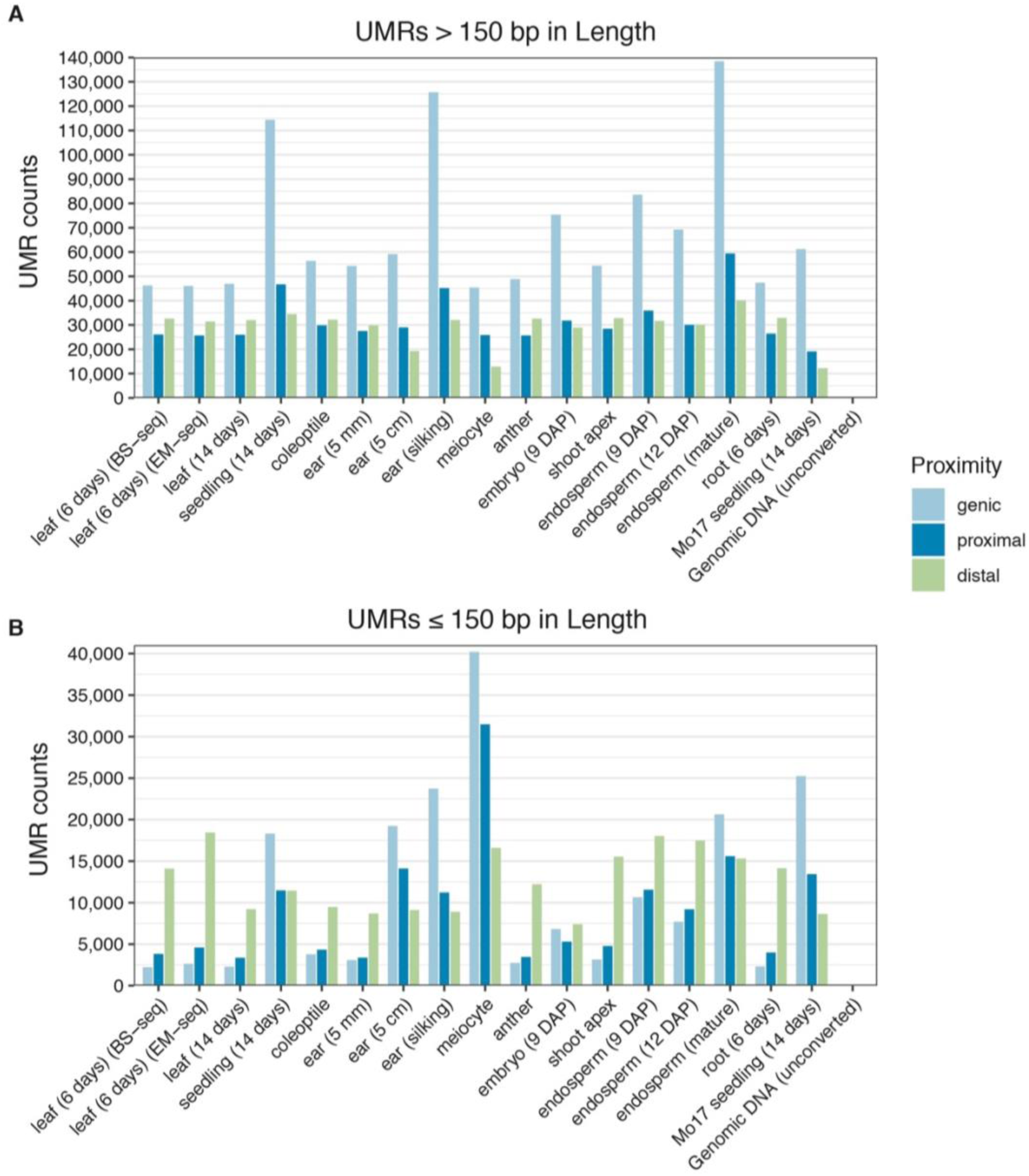
Counts of UMRs in different methylomes. Shown are counts of UMRs for each methylome, split by their proximities to AGPv4 annotated genes. Genic: overlaps genes by at least 1 bp. Proximal: within 5,000 bp of genes. Distal: > 5,000 bp from genes. UMRs are split by size into > 150 bp (**A**) and ≤ 150 bp (**B**).

**Supplementary Figure 3.5:**
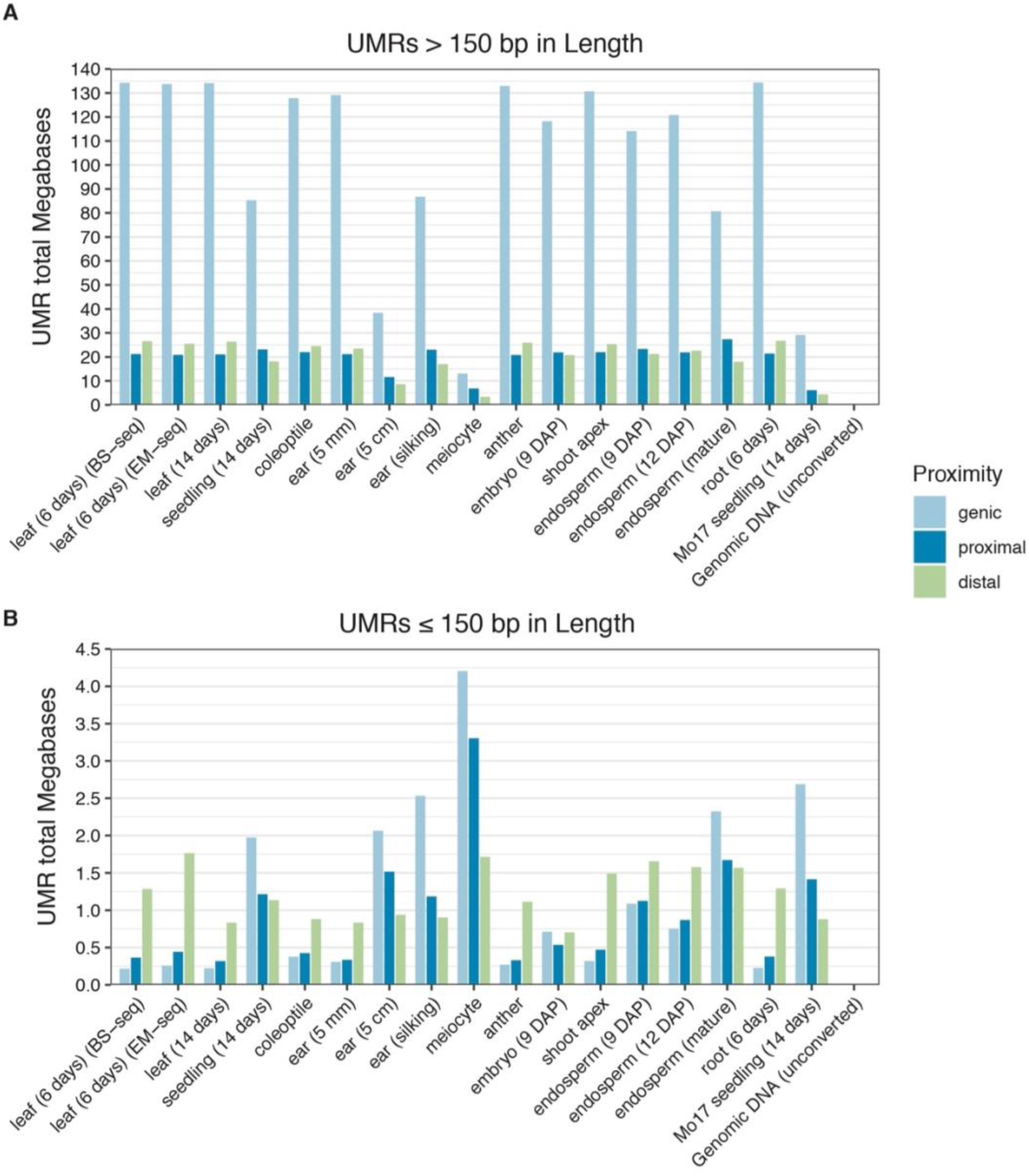
Total megabases of UMRs in different methylomes. Shown are sum of megabases of UMRs for each methylome, split by their proximities to AGPv4 annotated genes. Genic: overlaps genes by at least 1 bp. Proximal: within 5,000 bp of genes. Distal: > 5,000 bp from genes. UMRs are split by size into > 150 bp (**A**) and ≤ 150 bp (**B**).

**Supplementary Figure 3.6:**
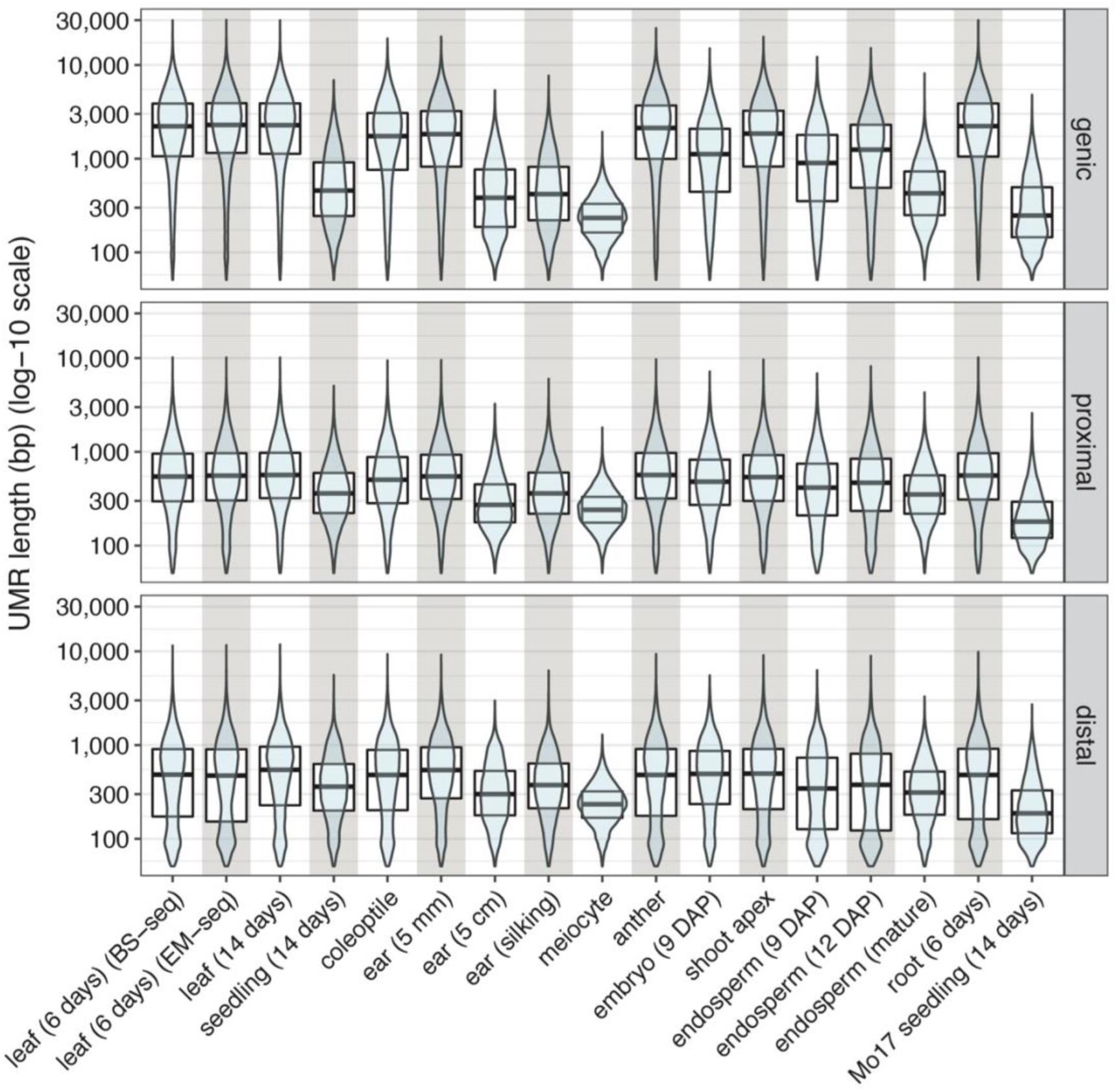
Length distributions of UMRs in different methylomes. Length distributions are shown as violin kernel density plots and boxplots containing quartiles. UMRs are split by their proximities to AGPv4 annotated genes. Genic: overlaps genes by at least 1 bp. Proximal: within 5,000 bp of genes. Distal: > 5,000 bp from genes. Apparent is the minimum UMR length cutoff of 50 bp.

**Supplementary Figure 3.7:**
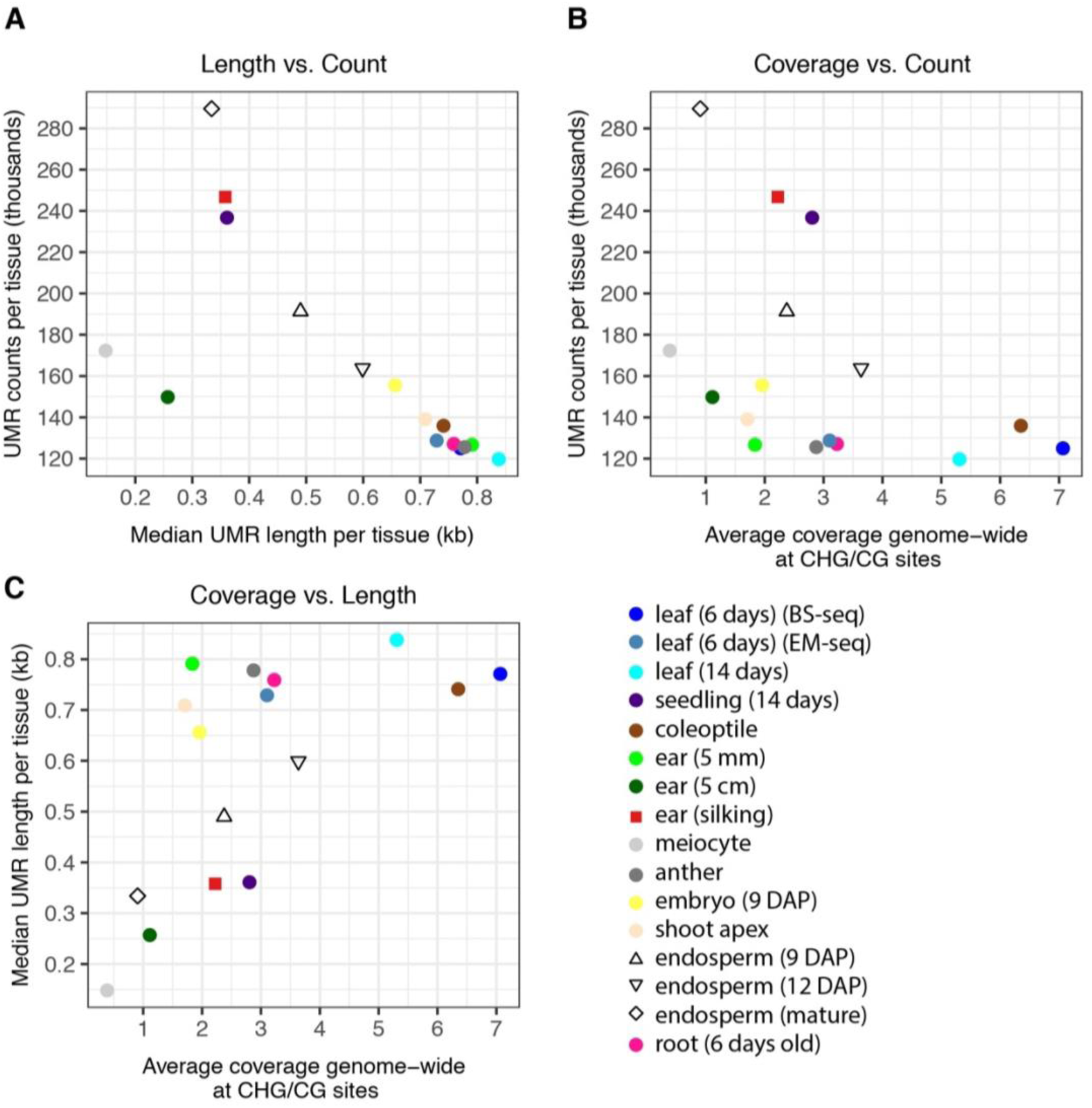
Relationships among UMR length, count, and coverage. Each point represents a single methylome (denoted in legend). **A:** UMR length vs. UMR count. **B:** UMR coverage vs. UMR count. **C:** UMR coverage vs. UMR length.

**Supplementary Figure 3.8:**
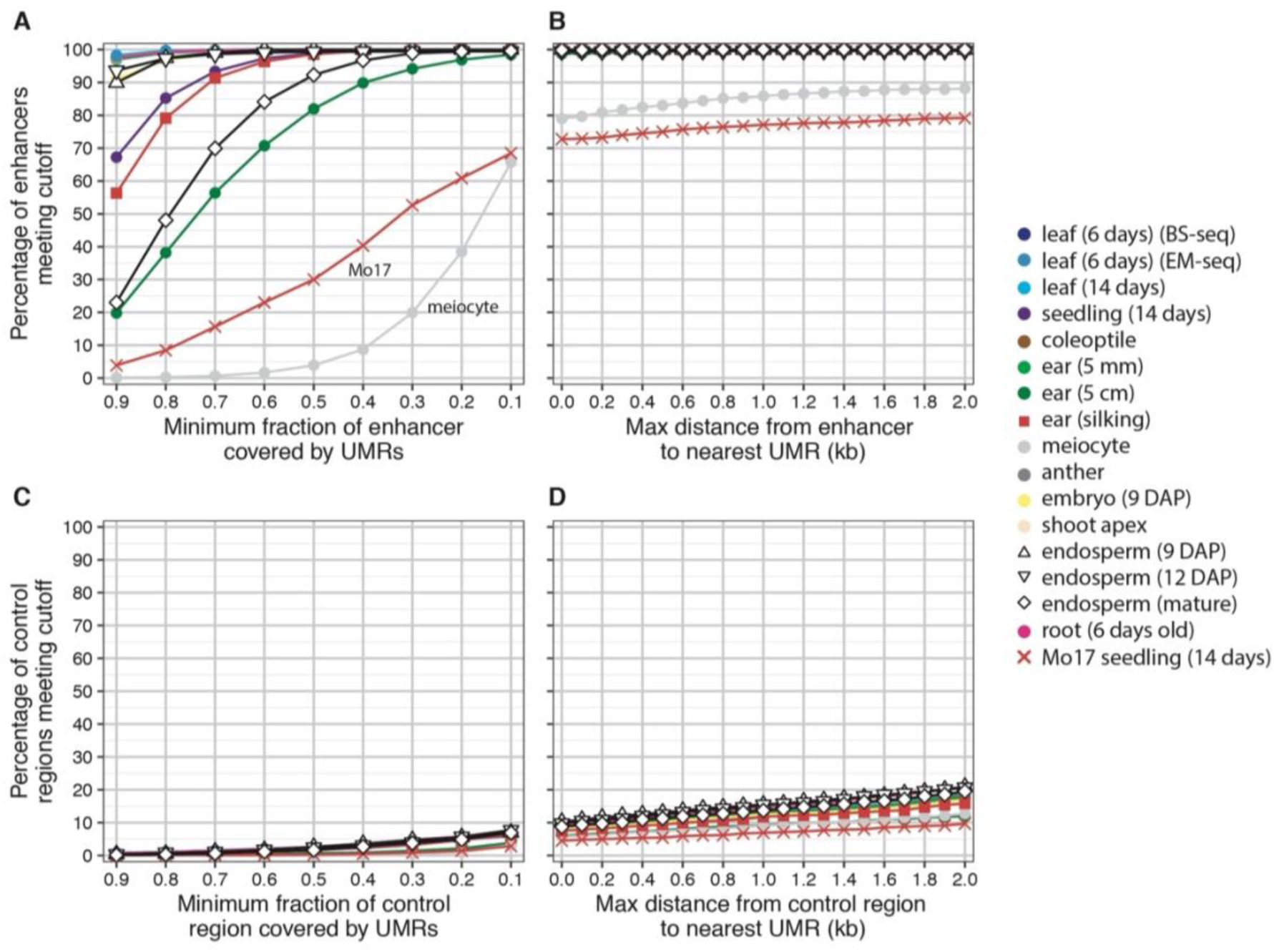
Overlap of UMRs with previously identified putative enhancers. Putative gene-distal enhancers from Oka, R., Zicola, J., Weber, B. *et al.* (2017), derived from husk and V2-IST tissues, were merged together. These enhancers were tested for overlap with the UMRs called in this manuscript. Each point and line represent a methylome (denoted on the right). **A & C:** The proportion of enhancers (**A**) and negative control regions (**C**) with different amounts of overlap with UMRs. For example, ∼30% of enhancers are covered by at least 50% of their lengths with UMRs derived the Mo17 seedling (14 days) methylome. **B & D:** The proportion of enhancers (**B**) and negative control regions (**D**) that have a UMR within the specified distance. Enhancers and control regions overlapping UMRs correspond to 0 kb.

**Supplementary Figure 3.9:**
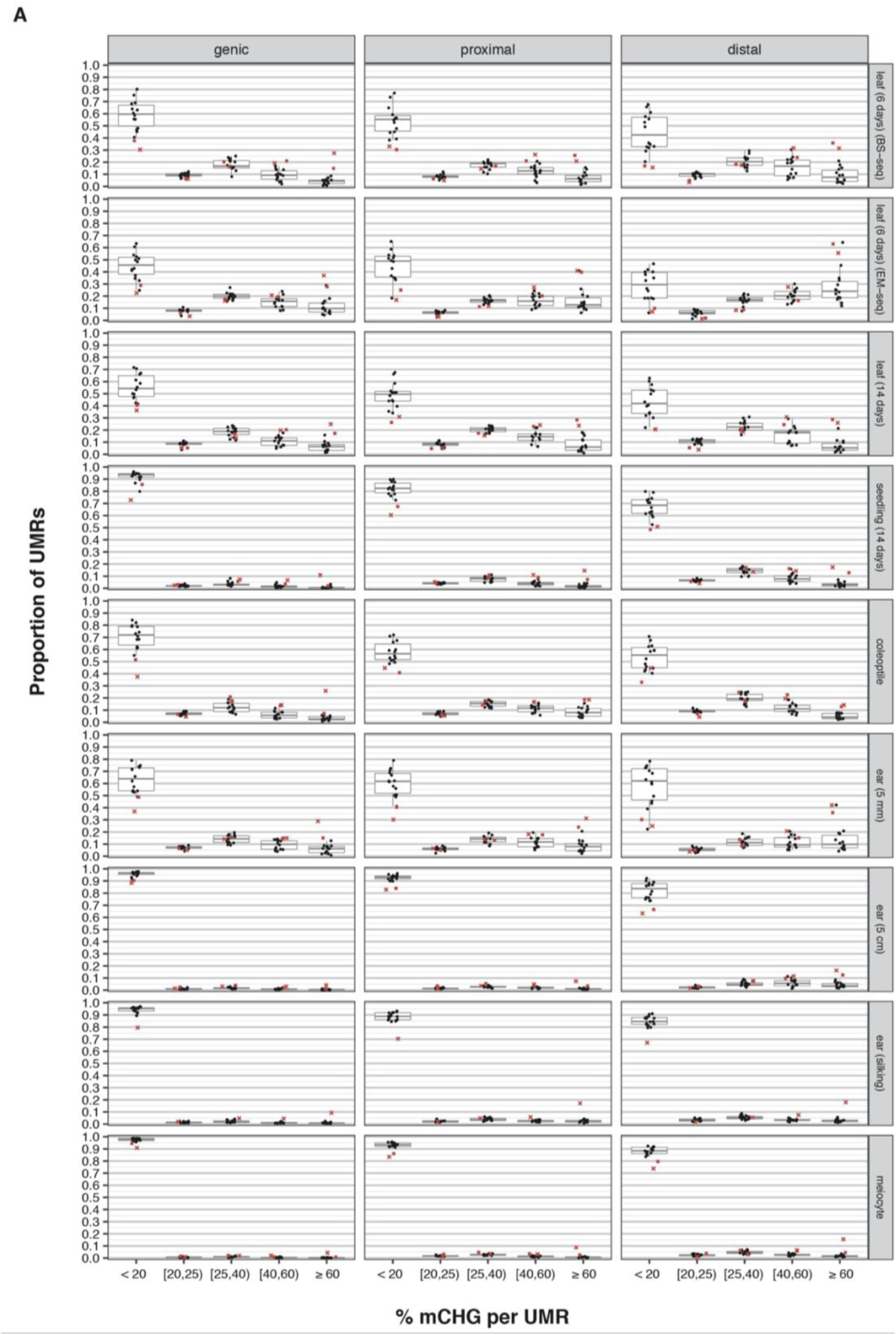
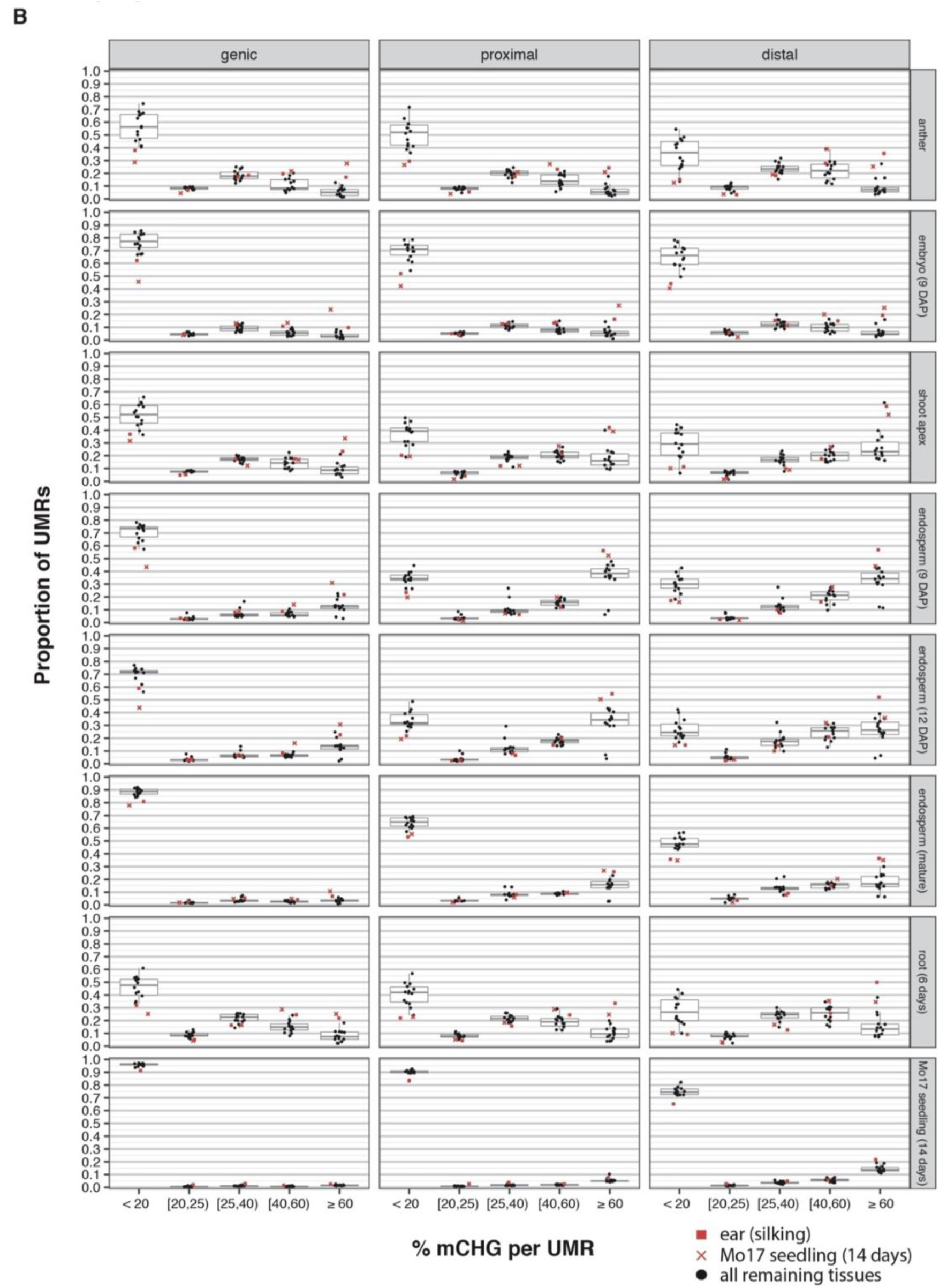
UMR (≤ 150 bp) stability across methylomes. Shown are the proportions of a methylome’s UMRs falling into discrete % mCHG bins in other methylomes. Each row corresponds to a set of UMRs (≤ 150 bp in length) identified in the primary methylome indicated on the right. Points within plots indicate the proportions of methylation levels in secondary methylomes. Boxplots indicate median and quartiles. Primary tissue UMRs are split by their proximities to AGPv4 annotated genes. Genic: overlaps genes by at least 1 bp. Proximal: within 5,000 bp of genes. Distal: > 5,000 bp from genes. Also shown are primary tissue UMRs that overlap constitutively transcribed exons. These serve as positive controls for developmentally stable UMRs. Ear (silking) and Mo17 seedling (14 days) are differentiated by shape and color (indicated in legend). **A & B:** Identical formatting but different primary methylomes.

**Supplementary Figure 3.10:**
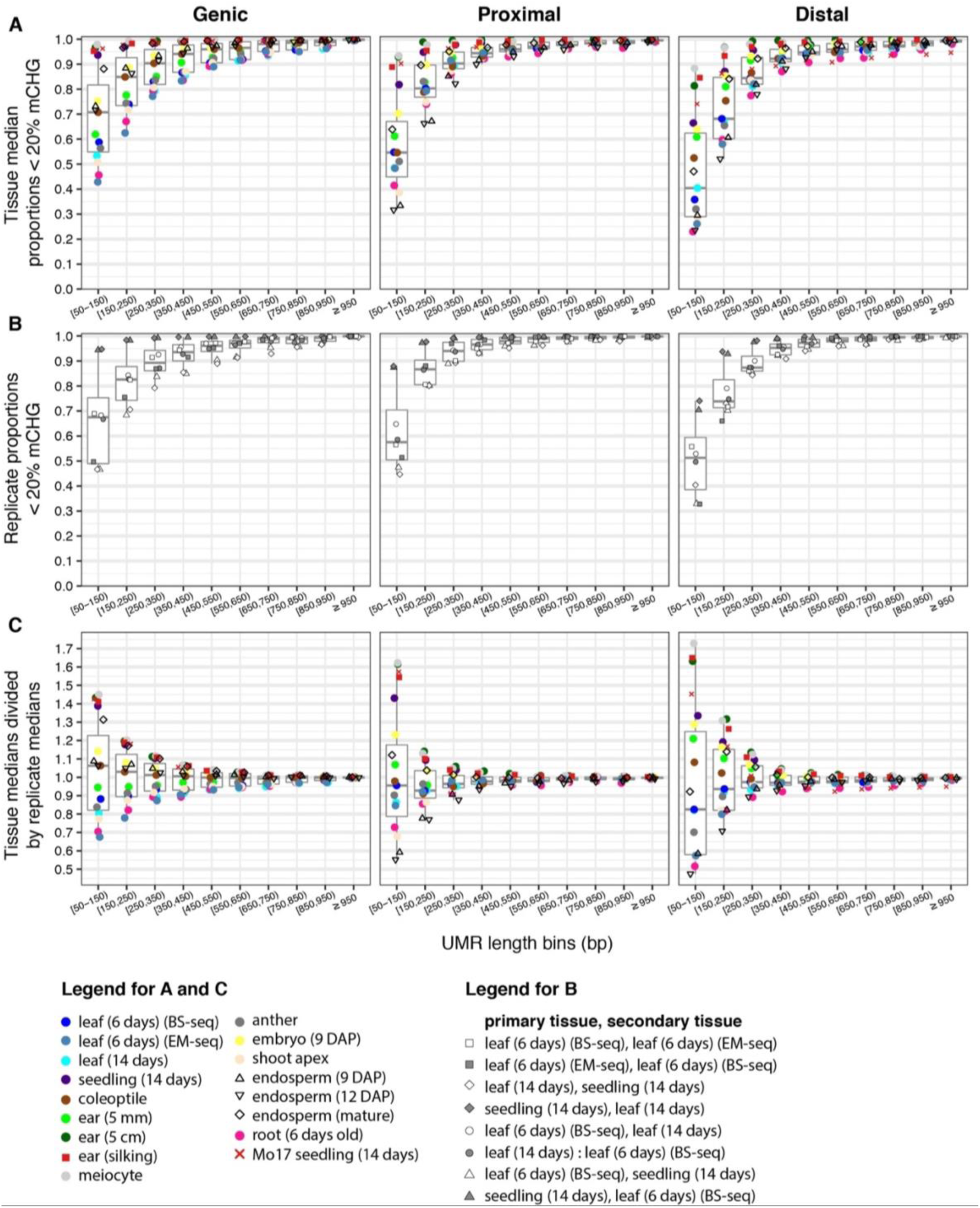
The stabilities of different length classes of UMRs. UMRs are split into discrete length bins (indicated on x-axes) and separated by their proximities to AGPv4 annotated genes. Genic: overlaps genes by at least 1 bp. Proximal: within 5,000 bp of genes. Distal: > 5,000 bp from genes. Each point corresponds to a single methylome (denoted at bottom). Boxplots indicate median and quartiles. **A:** Each point represents the median proportion of UMRs that remained below 20% in secondary tissues. **B:** Pairwise, reciprocal comparisons of samples that are either biological replicates or are developmentally similar tissues. Each point indicates the proportion of UMRs in the secondary tissue that remained < 20 % mCHG. **C:** The individual points in (**A**) divided by the medians indicated by boxplots in (**B**), but with ear (silking) and Mo17 seedling (14 days) removed from the median calculations. This figure serves to illustrate the discrepancy between replicate UMR stability and non-replicate UMR stability.

**Supplementary Figure 3.11:**
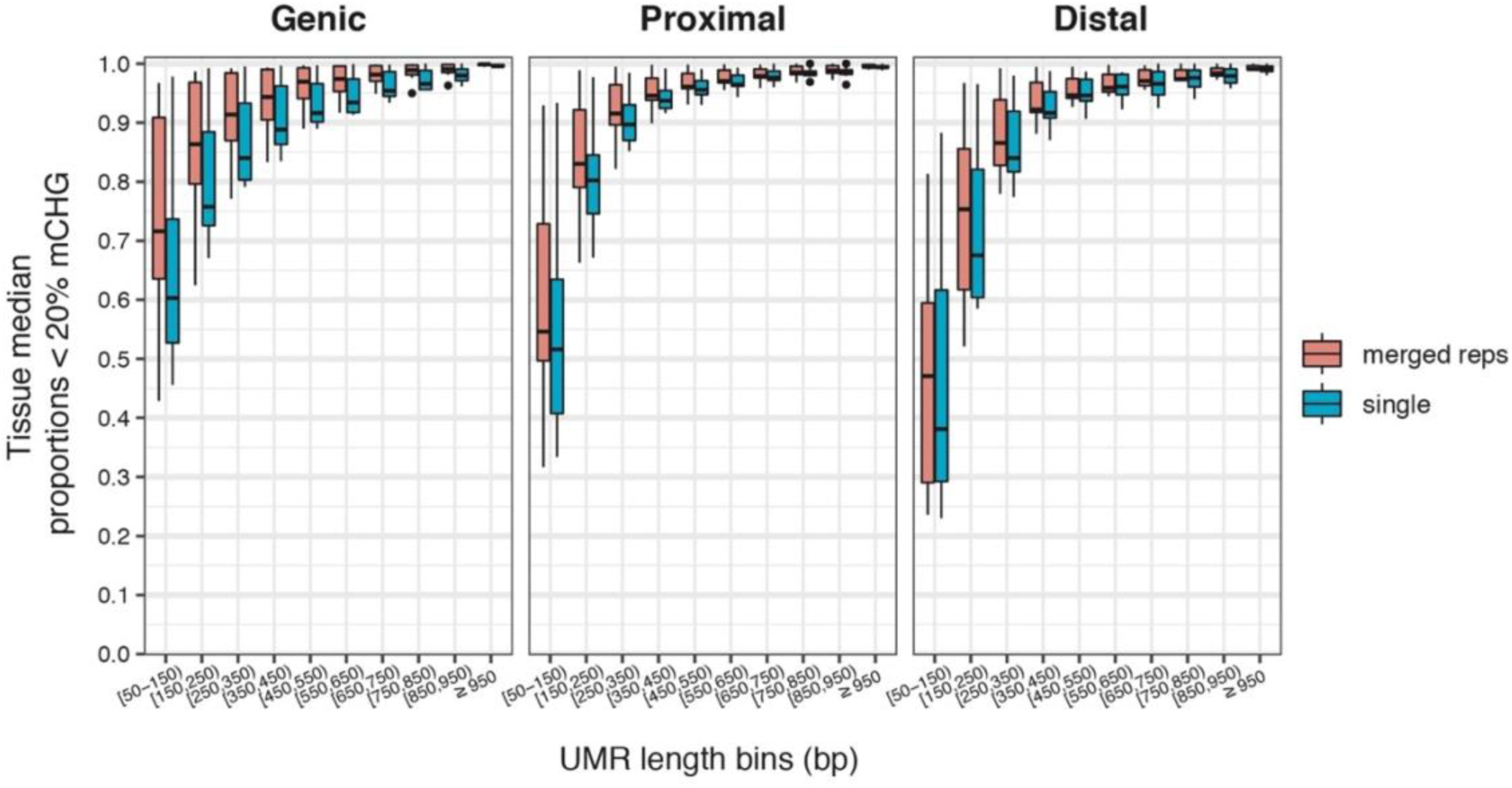
UMR stabilities separated by which methylomes comprise merged replicates. Shown are the same data as in sup. fig. 10A; however, ear (silking) and Mo17 seedling (14 days) methylomes are omitted and the datapoints are separated based on whether the methylome comprised merged biological replicates or a single biological sample (see sup. table 3.2). Boxplots indicate quartiles.

**Supplementary Figure 3.12:**
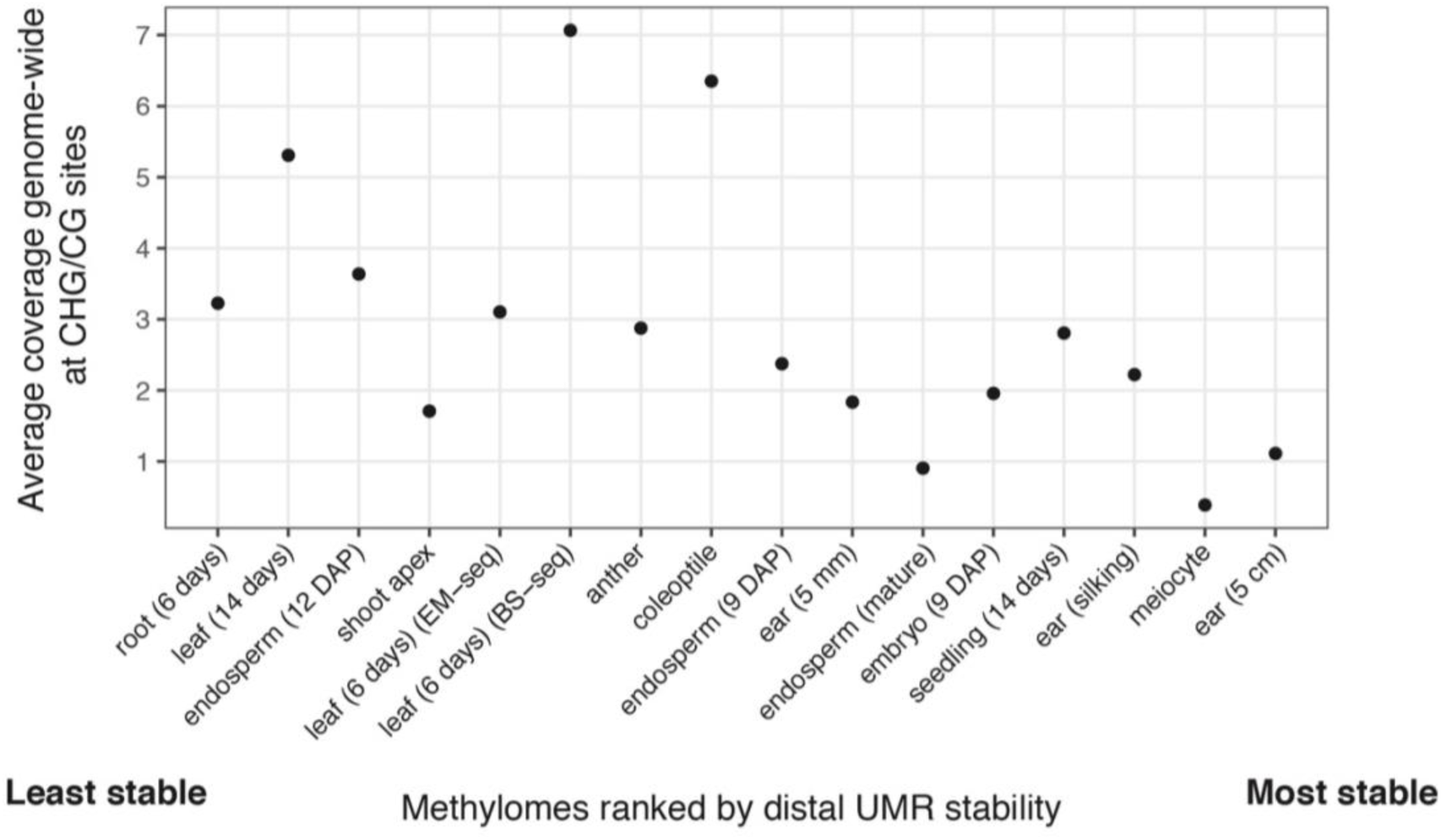
Methylomes ranked by distal UMR stability. Methylomes are ranked from least stable (left) to most stable (right), based on the rank sum of distal UMR baseline-corrected stabilities (i.e. stabilities shown in sup. fig. 3.10C). The genome-wide coverage is not restricted to UMRs.

**Supplementary Figure 3.13:**
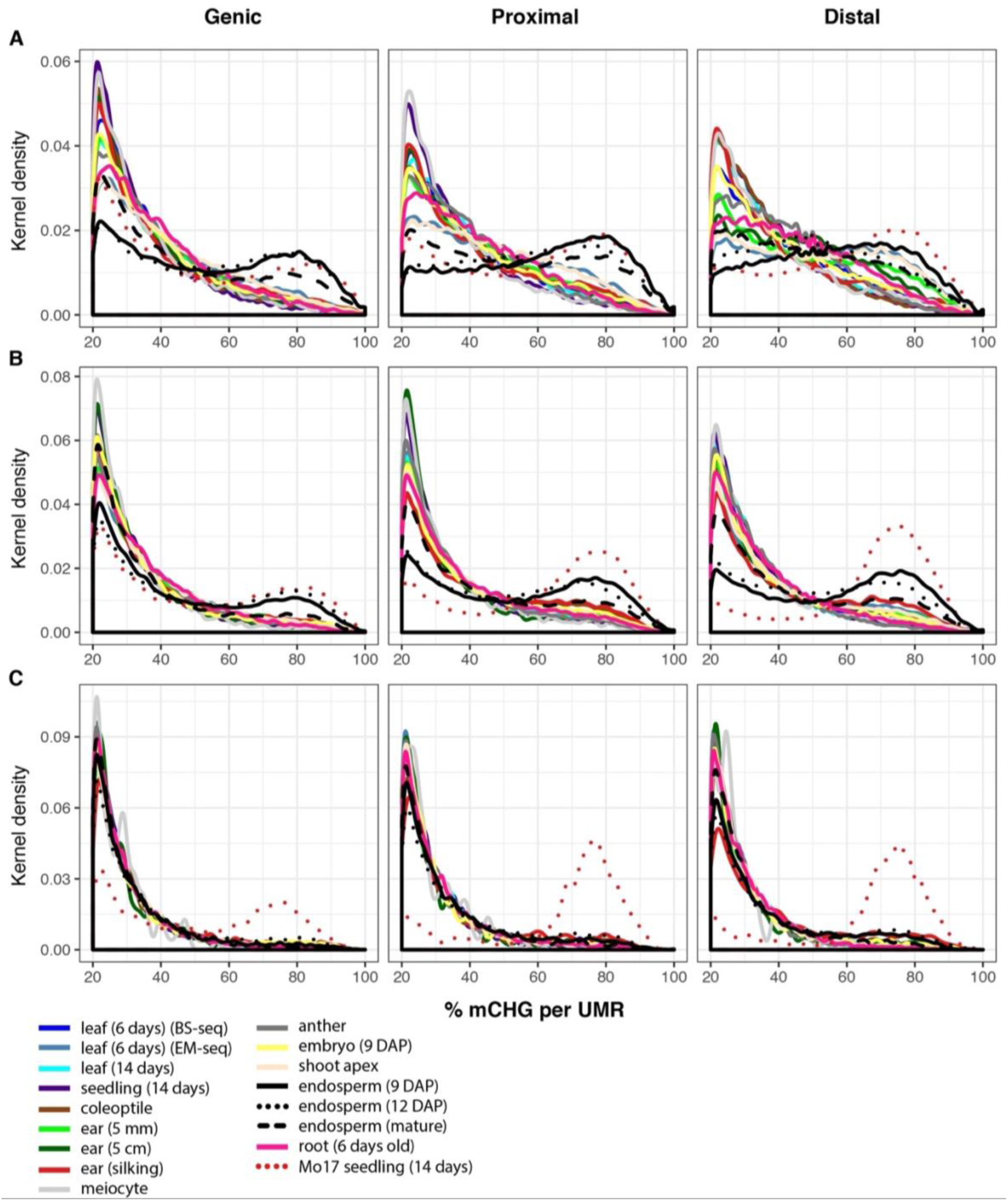
Methylation distributions of non-conserved UMRs. Shown here are frequency distributions (kernel density plots) of % mCHG of non-conserved UMRs. For each primary methylome’s UMR set (denoted in legend), all instances of UMRs exceeding 20 % mCHG in the secondary methylome are included in the density plots. All secondary methylome data are combined for each primary methylome in order to provide a single distribution. **A:** UMRs that are < 150 bp in length. **B:** UMRs 150-500 bp. **C:** UMRs > 500 bp.

**Supplementary Figure 3.14:**
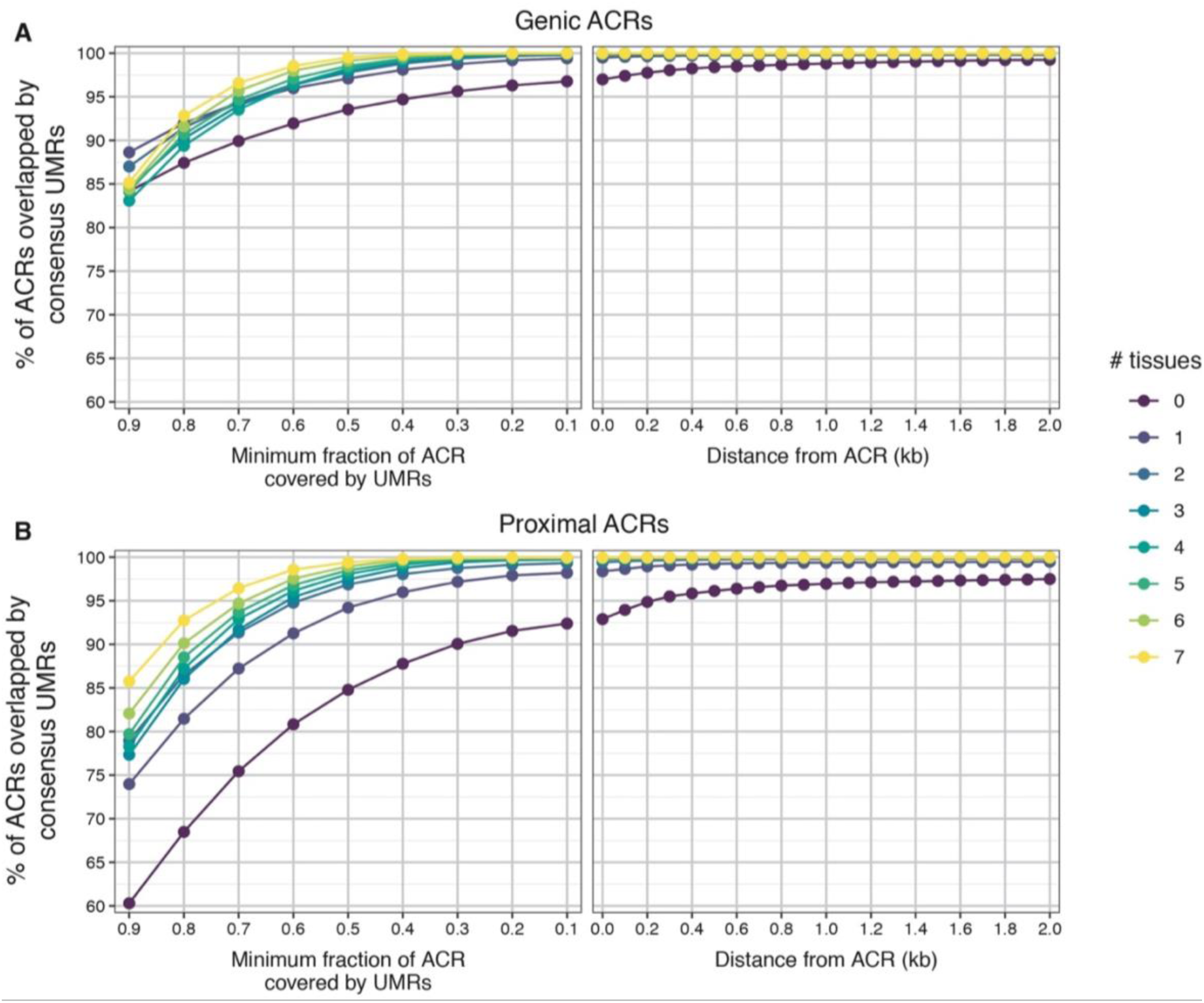
Genic and proximal ACRs overlapping UMRs. Shown are the fractional and distance overlaps akin to those shown in fig. 3.4B. # tissues denotes the ACR overlap category; e.g. category 0 ACRs overlap no ACRs from other tissues.

**Supplementary Figure 3.15:**
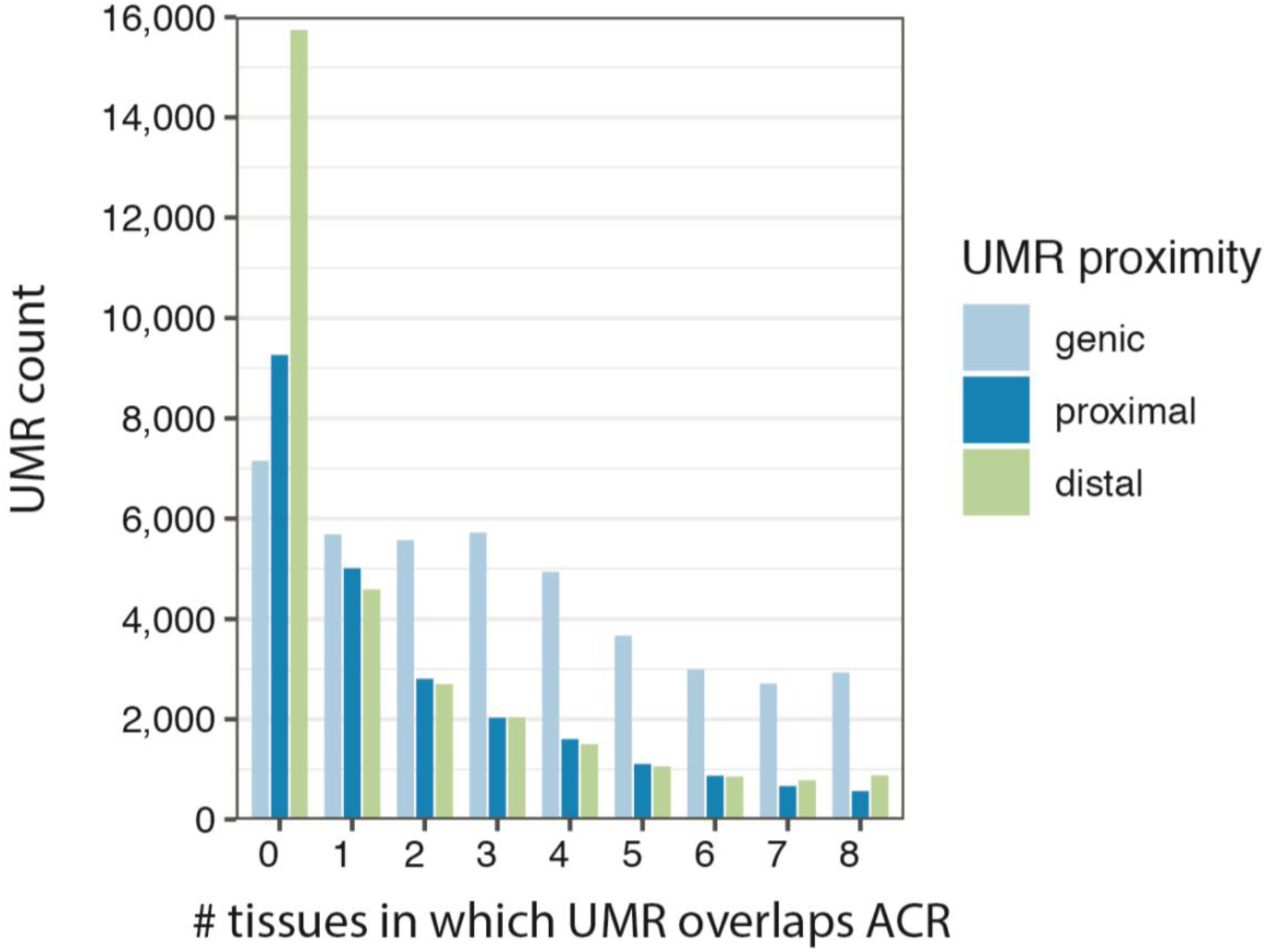
Counts of UMR and ACR overlaps. Shown are counts from the consensus UMR set. Overlaps are ≥ 1 bp. 0 on the x-axis denotes UMRs that overlap no ACRs. 8 on the x-axis denotes UMRs that overlap ACRs in all 8 tissues.

**Supplementary Figure 3.16:**
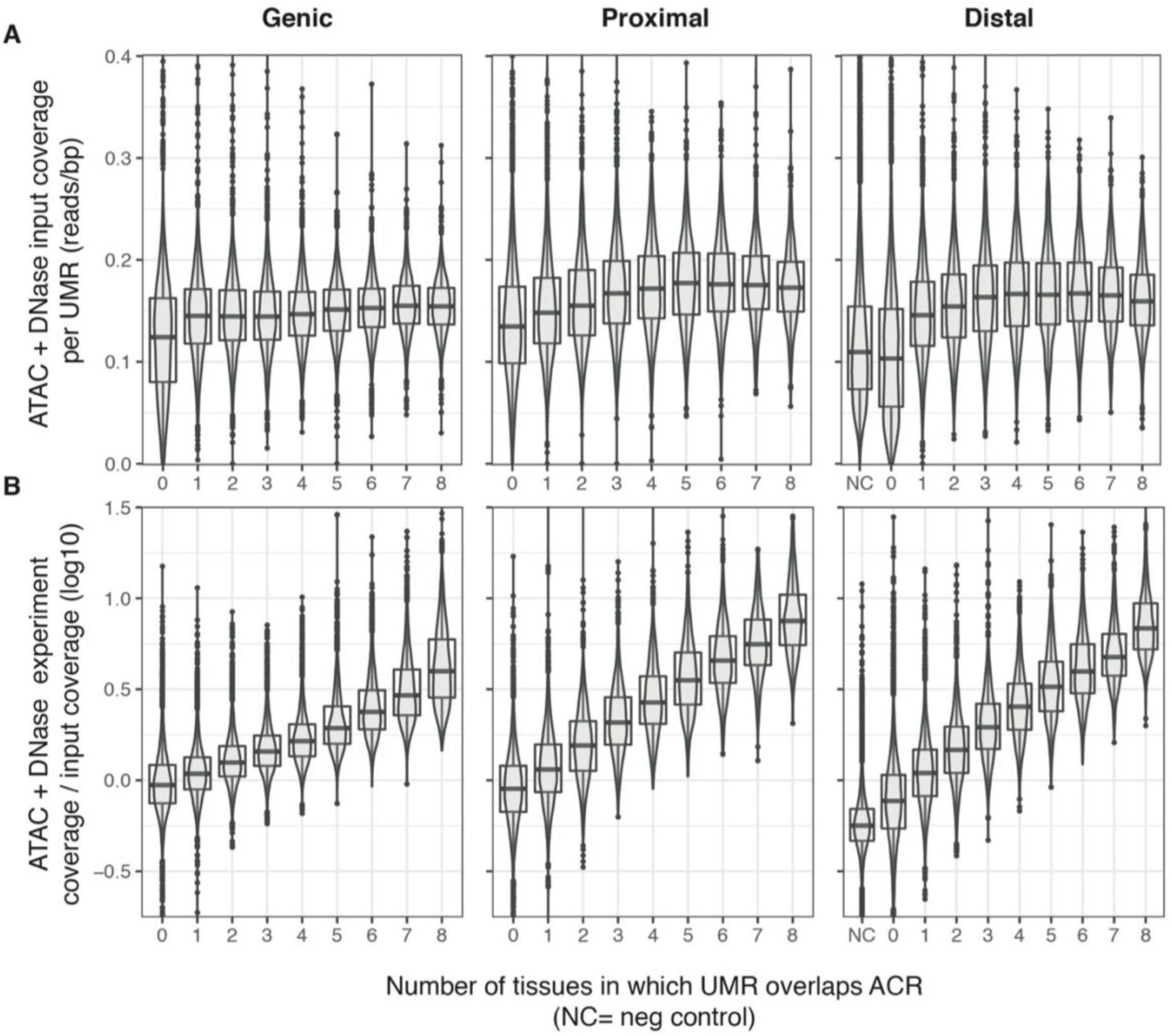
ATAC and DNase coverages at UMRs. ATAC-seq reads (ear (5 mm), leaf (6 day), and mesophyll) and DNase-seq reads (inner husk and inner stem) were pooled together, along with their respective input controls, to get aggregate read counts at each UMR. UMRs are separate into subcategories based on the number of tissues in which a UMR overlaps an ACR. The category 0 means that the UMR never overlaps ACRs. NC: distal UMR negative control regions. **A:** The coverages of the input reads. Input read counts per UMR are divided by the length of the UMR. **B:** The experimental read counts per UMR divided by the input read counts per UMR. Boxplots denote quartiles. Data points denote outliers.

**Supplementary Figure 3.17:**
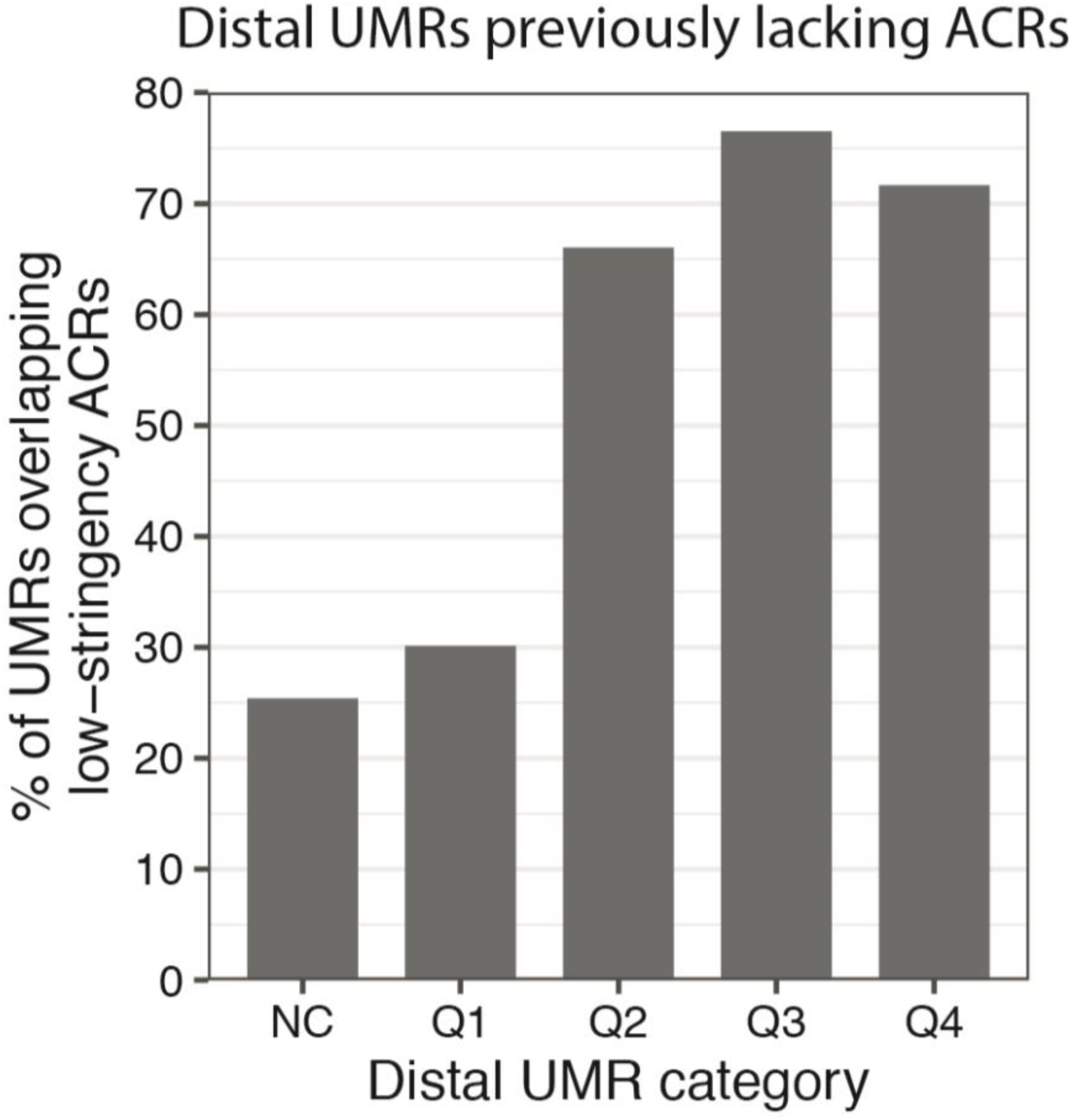
UMRs overlapping relaxed-stringency ACR peaks. For the distal UMRs previously lacking any ACR overlaps, the UMRs were split into quartiles Q1-Q4. Genrich peak-calling stringency was relaxed and the UMRs re-analyzed for overlaps with the new ACR peaks. The percentages on the y-axis show what percent of each UMR group now overlaps the new ACR peaks.

**Supplementary table 3.2.** A summary of UMRs called in various maize tissues.

| sample | reps single or merged | mapped genome | CHG/CG average coverage | median UMR length (bp) | Counts, UMRs below 150 bp |  |  | Counts, UMRs above 150 bp |  |  | Total Mb, UMRs below 150 bp |  |  | Total Mb, UMRs above 150 bp |  |  |
| --- | --- | --- | --- | --- | --- | --- | --- | --- | --- | --- | --- | --- | --- | --- | --- | --- |
|  |  |  |  |  | genic | proximal | distal | genic | proximal | distal | genic | proximal | distal | genic | proximal | distal |
| endosperm (12 DAP) | merge | AGPv4 | 3.64 | 599 | 7,686 | 9,198 | 17,483 | 69,204 | 30,056 | 30,067 | 0.8 | 0.9 | 1.6 | 120.8 | 21.9 | 22.6 |
| leaf (14 days) | single | AGPv4 | 5.31 | 838 | 2,271 | 3,354 | 9,209 | 46,884 | 25,945 | 32,019 | 0.2 | 0.3 | 0.8 | 134.1 | 21.0 | 26.4 |
| seedling (14 days) | merge | AGPv4 | 2.81 | 361 | 18,310 | 11,467 | 11,429 | 114,362 | 46,684 | 34,472 | 2.0 | 1.2 | 1.1 | 85.3 | 23.1 | 18.1 |
| ear (5 cm) | merge | AGPv4 | 1.11 | 257 | 19,219 | 14,100 | 9,104 | 59,138 | 28,994 | 19,247 | 2.1 | 1.5 | 0.9 | 38.4 | 11.6 | 8.6 |
| ear (5 mm) | single | AGPv4 | 1.83 | 791 | 3,068 | 3,374 | 8,674 | 54,242 | 27,493 | 29,834 | 0.3 | 0.3 | 0.8 | 129.1 | 21.1 | 23.5 |
| leaf (6 days) (BS-seq) | single | AGPv4 | 7.07 | 771 | 2,221 | 3,809 | 14,092 | 46,201 | 26,045 | 32,611 | 0.2 | 0.4 | 1.3 | 134.2 | 21.2 | 26.5 |
| root (6 days) | single | AGPv4 | 3.23 | 759 | 2,312 | 3,988 | 14,151 | 47,336 | 26,404 | 32,903 | 0.2 | 0.4 | 1.3 | 134.3 | 21.4 | 26.7 |
| embryo (9 DAP) | single | AGPv4 | 1.96 | 656 | 6,822 | 5,300 | 7,384 | 75,361 | 31,790 | 28,879 | 0.7 | 0.5 | 0.7 | 118.2 | 21.9 | 20.7 |
| endosperm (9 DAP) | single | AGPv4 | 2.38 | 490 | 10,635 | 11,560 | 18,035 | 83,566 | 35,897 | 31,641 | 1.1 | 1.1 | 1.7 | 114.1 | 23.3 | 21.2 |
| anther | merge | AGPv4 | 2.88 | 778 | 2,753 | 3,461 | 12,218 | 48,847 | 25,675 | 32,612 | 0.3 | 0.3 | 1.1 | 132.9 | 20.8 | 25.9 |
| shoot apex | single | AGPv4 | 1.71 | 709 | 3,148 | 4,777 | 15,532 | 54,347 | 28,418 | 32,807 | 0.3 | 0.5 | 1.5 | 130.7 | 22.0 | 25.3 |
| coleoptile | merge | AGPv4 | 6.35 | 741 | 3,768 | 4,332 | 9,475 | 56,321 | 29,836 | 32,180 | 0.4 | 0.4 | 0.9 | 127.8 | 22.0 | 24.5 |
| endosperm (mature) | merge | AGPv4 | 0.91 | 334 | 20,628 | 15,601 | 15,315 | 138,458 | 59,395 | 40,131 | 2.3 | 1.7 | 1.6 | 80.7 | 27.3 | 17.9 |
| meiocyte | single | AGPv4 | 0.39 | 148 | 40,187 | 31,476 | 16,591 | 45,378 | 25,845 | 12,828 | 4.2 | 3.3 | 1.7 | 13.0 | 6.8 | 3.4 |
| ear (silking) | single | AGPv4 | 2.22 | 358 | 23,733 | 11,212 | 8,900 | 125,692 | 45,162 | 32,045 | 2.5 | 1.2 | 0.9 | 86.7 | 23.0 | 17.0 |
| Mo17 seedling (14 days) | single | AGPv4 | 1.02 | 206 | 25,235 | 13,432 | 8,630 | 61,189 | 19,114 | 12,219 | 2.7 | 1.4 | 0.9 | 29.2 | 6.1 | 4.3 |
| leaf (6 days) (EM-seq) | merge | AGPv4 | 3.10 | 729 | 2,629 | 4,599 | 18,442 | 46,039 | 25,629 | 31,407 | 0.3 | 0.4 | 1.8 | 133.7 | 20.8 | 25.5 |

**Supplementary Table 3.4:** Estimating false negative rates with a transcription start site positive control. The transcription starts sites (+/-500 bp) from 307 genes that are constitutively highly expressed were tested for overlap frequency with ACRs.

| % of TSS overlapped by ACR | % of TSS +/- 500 bp overlapped by ACR | Tissue | peak calling method | Experimental method | # genic peaks | # proximal peaks | # distal peaks |
| --- | --- | --- | --- | --- | --- | --- | --- |
| 34.5 | 36.8 | ear (5 mm) | Genrich | ATAC-seq | 5,781 | 5,750 | 2,261 |
| 49.2 | 59.3 | ear (5 mm) | Called by Ricci et al. (2019) | ATAC-seq | 9,745 | 8,213 | 9,330 |
| 59.3 | 60.6 | leaf (6 day) | Genrich | ATAC-seq | 9,534 | 9,156 | 3,427 |
| 67.8 | 75.2 | leaf (6 day) | Called by Ricci et al. (2019) | ATAC-seq | 12,587 | 9,243 | 10,651 |
| 80.1 | 88.9 | ear (1 cm) | Genrich | Differential Mnase | 32,041 | 34,488 | 11,965 |
| 38.1 | 42.7 | coleoptilar node | Genrich | Differential Mnase | 4,529 | 7,331 | 2,818 |
| 67.4 | 75.6 | root tip | Genrich | Differential Mnase | 17,403 | 18,823 | 6,756 |
| 58.0 | 61.6 | mesophyll | Genrich | ATAC-seq | 14,763 | 21,707 | 7,484 |
| 70.4 | 74.9 | inner husk | Genrich | Dnase-seq | 29,318 | 26,315 | 10,210 |
| 81.8 | 88.3 | inner stem | Genrich | Dnase-seq | 59,219 | 28,718 | 10,831 |

